# Unraveling systemic responses to NQO1-activated IB-DNQ and Rucaparib single and dual agent therapy in triple-negative breast cancers

**DOI:** 10.1101/2024.05.15.594427

**Authors:** Avery M. Runnebohm, H.R. Sagara Wijeratne, Sarah A. Peck Justice, Aruna B. Wijeratne, Gitanjali Roy, Jarrett J. Smith, Naveen Singh, Paul Hergenrother, David A. Boothman, Edward A. Motea, Amber L. Mosley

## Abstract

Triple negative breast cancer (TNBC) is a highly aggressive breast cancer that is unresponsive to hormonal therapies. One potential TNBC-specific therapeutic target is NQO1, as it is highly expressed in many TNBC patients and lowly expressed in non-cancer tissues. Here we use a derivative of DNQ, isobutyl-deoxynyboquinone (IB-DNQ) that is more potent and specific in killing TNBC cells than NQO1-activator β-lapachone while displaying strong NQO1-dependence. We evaluated the cellular signaling changes that occur following 4-hour treatment of TNBC cells with either single agent or combination IB-DNQ and / or PARP inhibitor (Rucaparib). Short treatments (4 hours) with IB-DNQ alone or combined with the PARP inhibitor Rucaparib revealed few changes in protein abundance but significant rapid alterations in protein phosphorylation and thermal stability, with clear synergy in the combination treatment. Key phosphorylated targets linked to RNA Polymerase II inhibition and DNA damage response were altered during our short time treatment. Thermal proteome profiling (TPP) identified novel, combination-specific changes in protein biophysical state suggesting new therapeutic vulnerabilities in TNBC cells. Our findings highlight how even brief treatments can uncover distinct biophysical protein changes via TPP, offering a resource for mechanistic studies of IB-DNQ mechanism of action and the development of NQO1-activated therapeutics for TNBC treatment.

## BACKGROUND

Triple negative breast cancer (TNBC) is a highly aggressive form of breast cancer that accounts for 15-25% of cases (1). TNBC is more aggressive than most other breast cancer types, with approximately 46% of patients developing metastases and an average tumor recurrence of 1.2 years (1–3). TNBC is characterized by a lack of estrogen, progesterone, and human epidermal growth factor 2 receptors, which render these tumors unresponsive to commonly used hormonal therapies (4,5). Conventional treatments of TNBC include alkylating agents, anti-microtubule agents, anti-metabolites, and platinum (2). For advanced cases of TNBC, new immunotherapies have emerged, but even with these therapies, TNBC has a 5-year survival rate of 8-16% (2,3). An efficacious tumor type-specific treatment would greatly improve outcomes for patients with TNBC.

NAD(P)H:quinone oxidoreductase 1 (NQO1) bioactivatable agents have been shown previously to cause tumor-selective DNA damage and cell death (6,7). The NQO1 bioactivatable drug β-lapachone (ARQ761 in clinical form) has been tested in clinical trials for patients with solid tumors with metastasis or that have recurrent or nonresectable tumors (8). However, two other NQO1 bioactivatable drugs, deoxynyboquinone (DNQ) and its derivative isobutyl deoxynyboquinone (IB-DNQ), have recently been shown to be more potent than β-lapachone in killing cancer cells, with one report demonstrating 60-fold higher potency of IB-DNQ compared to β-lapachone in TNBC cells (9–11). β-lapachone has been shown to induce DNA damage and hyperactivation of the DNA repair enzyme PARP1, leading to reduction of essential nucleotides and programmed necrosis (12,13). Combination treatment of TNBC with sub-lethal concentrations of β-lapachone and PARP inhibition has been shown to kill cells through apoptosis rather than programmed necrosis, suggesting that tumor-selective NQO1 activators may have multiple paths that can be exploited for cancer treatment. For this reason, establishing a full understanding of NQO1 activator mechanism of action is imperative to unlock multiple potential treatment avenues for aggressive diseases such as TNBC.

These drugs are promising in the treatment of TNBC and other solid tumors with overexpression of NQO1 and an elevated NQO1:catalase expression ratio (14). Many solid tumors, including those from thyroid, adrenal, breast, ovarian, colon, cornea, and non-small cell lung cancer tissues, have significantly increased NQO1 expression compared to normal tissues suggesting broad potential utility for NQO1 inhibition in cancer treatment (15). NQO1 is a quinone reductase that catalyzes two-electron reduction of quinones to hydroquinones using NADH or NADPH as electron donors (16,17). NQO1 bioactivatable drugs, including β-lapachone and IB-DNQ, undergo oxidoreduction by NQO1 which results in unstable hydroquinones that spontaneously convert back to the original compound (13) (Figure 1a). This NQO1-dependent futile redox cycling oxidizes NAD(P)H to create reactive oxygen species (ROS), most of which are superoxide radicals (18). Superoxide dismutase converts superoxide radicals into oxygen and hydrogen peroxide, and catalase catalyzes degradation of hydrogen peroxide to protect cells from damage (reviewed in (19)). Treating tumors that highly express NQO1 and lowly express catalase with NQO1 bioactivatable drugs has been shown to cause ROS induced-DNA damage specifically within cancer cells. Therefore, NQO1 selective agents including β-lapachone, DNQ, and IB-DNQ have potential to be used as single agent treatments, but strategies to improve efficacy while minimizing toxicity are needed and could provide avenues for rapid combination therapies if cancer cell resistance occurs over time.

**Figure 1:**
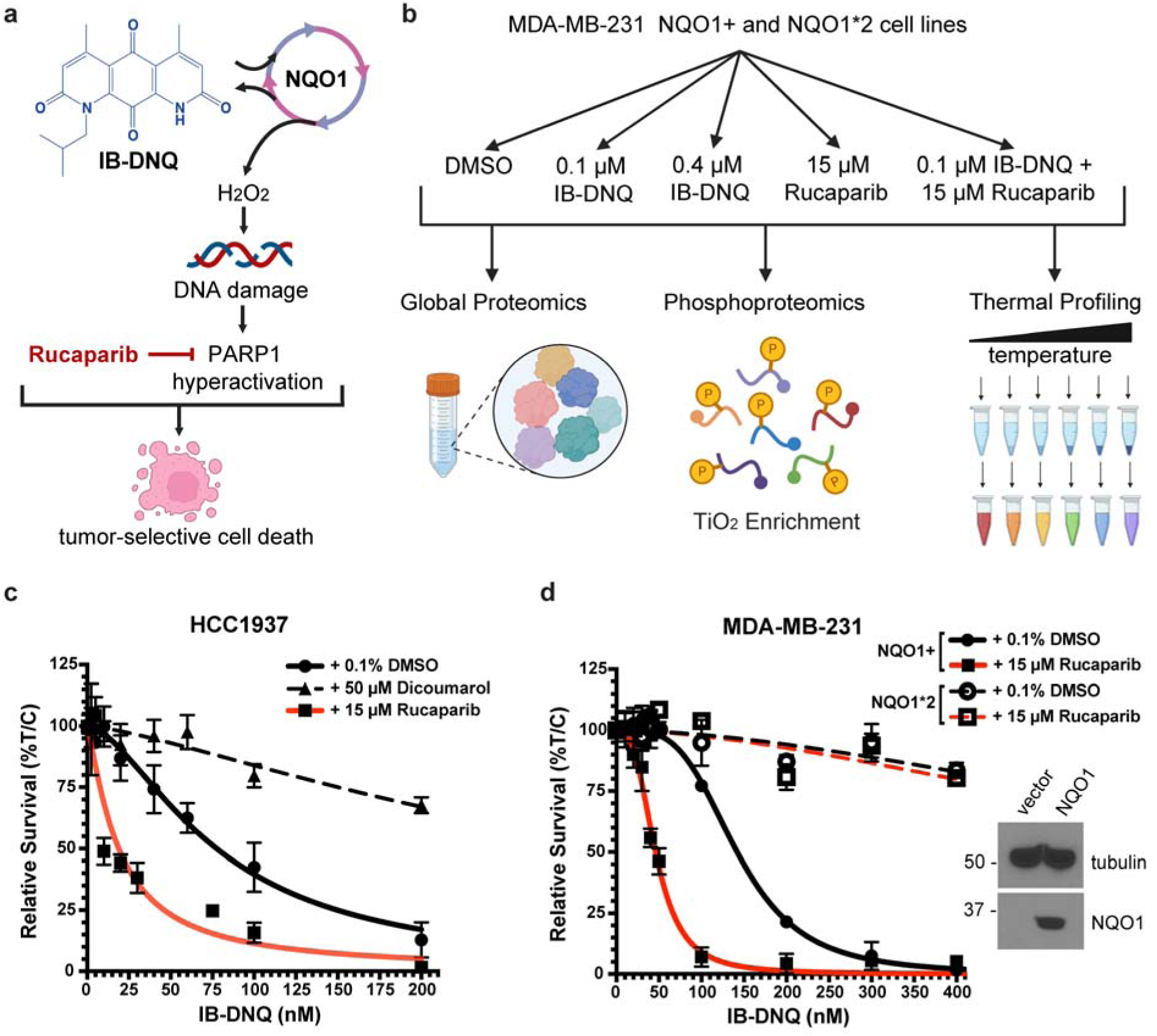
Treatment with the bioactivatable drug IB-DNQ in combination with PARP1 inhibitor, Rucaparib, leads to tumor-selective cell death. a) IB-DNQ is a futile-cycling substrate of NQO1. IB-DNQ processing by NQO1 produces reactive oxygen species resulting in DNA damage. Inhibition of PARP1-mediated DNA repair mechanisms via Rucaparib in combination with IB-DNQ treatment leads to tumor-selective cell death. b) For proteomics experiments, NQO1+ and NQO1*2 MDA-MB-231 cells were treated with DMSO, 0.1 µM (sublethal) or 0.4 µM (lethal) IB-DNQ, 15 µM Rucaparib, or a combination treatment of 0.1 µM IB-DNQ and 15 µM Rucaparib. Samples were analyzed using global proteomics, phosphoproteomics, and TPP. c) HCC1937 cells (primary ductal carcinoma, NQO1+) were treated with increasing doses of IB-DNQ in combination with DMSO, Dicoumarol (NQO1 inhibitor), or Rucaparib (PARP1 inhibitor). Cell survival at 7-days post 4-hour treatment is plotted for increasing IB-DNQ concentrations. Cells co-treated with Dicoumarol are represented by the dashed black line, and cells co-treated with Rucaparib are shown by the red line. d) MDA-MB-231 triple negative breast cancer cells, which express the NQO1*2 variant (rapidly degraded NQO1), were altered with a vector to stably overexpress wild-type NQO1 (NQO1+) or an empty vector control. Cells were treated with increasing doses of IB-DNQ in combination with DMSO or 15 µM Rucaparib. Cell survival 7-days post 4-hour treatment is plotted for increasing IB-DNQ concentrations. NQO1*2 cells are represented by dashed lines, and NQO1+ cells are represented by solid lines. Black lines show co-treatment with DMSO, and red lines show co-treatment with Rucaparib. %T/C = % of treated / control cells. Western blot insets show abundance of NQO1 in NQO1+ and vector containing MDA-MB-231 cells. Tubulin was used as the loading control.

DNA repair mechanisms have long been targets for cancer therapies because of the common vulnerability of many cancer cells to DNA damage (20,21). Poly(ADP-ribose) polymerase-1 (PARP1) is a poly-ADP-ribosyltransferase that mediates poly-ADP-ribosylation (PARylation) and plays an important role in multiple DNA repair pathways (22–24). PARP1 inhibitors (PARPi), including Olaparib, Rucaparib, Niraparib, and Talazoparib, have been approved by the FDA for treatment of ovarian and breast cancers (22), but unfortunately are often not effective as a single agent treatment in TNBCs due to PARPi resistance mechanisms that develop in TNBC (25). Huang et al (2016) showed that pretreating NQO1+ TNBC cells with a non-lethal concentration of Rucaparib, followed by co-treatment with β-lapachone leads to tumor-selective cell death (12). PARP1 becomes hyperactive following treatment with β-lapachone, where self PARylation (PAR-PARP1) inactivates the protein and prevents PARP1 from binding DNA and activating DNA repair processes (26,27). Therefore, inhibiting PARP1 with Rucaparib prior to NQO1 inhibition enhances DNA damage and apoptosis while lowering the efficacious dose of NQO1 activators and decreasing toxicity (Figure 1a) (12). To date, PARP inhibition has not been studied in combination with next generation NQO1 substrate-based activators such as IB-DNQ. Additionally, the mechanisms of NQO1-dependent TNBC cellular responses, which could form the basis for cancer cell development of resistance, have not been investigated with broad-focused multiomics analyses.

The precise molecular mechanisms leading to tumor-selective cell death by dual agent treatment with NQO1 bioactivatable drug IB-DNQ and its potential for combination treatment with Rucaparib have not been studied. Here we are using several discovery proteomics techniques (e.g. global proteome abundance analysis, phosphoproteomics, and thermal proteome profiling) to gain a better understanding of the effects of IB-DNQ and Rucaparib on protein abundance, protein phosphorylation, and protein thermal stability in TNBC cells with elevated NQO1 protein levels (Figure 1b). We report that 4-hour treatment with IB-DNQ and Rucaparib results in significant changes in protein phosphorylation and thermal stability despite minimal impacts on protein abundance. Deeper analysis into the proteins with changes in phosphorylation and / or thermal stability revealed alterations in the DNA damage response, RNA Polymerase II transcription, chromatin modifications, and cell cycle related proteins. Specifically, we observe changes in phosphorylation of H2AX and in protein-complex stability of chromatin remodeling machinery, indicative of persistent DNA damage following 4-hour treatment. We show significant changes in cyclin dependent kinase activity and protein thermal stability of several cell cycle, DNA repair, and chromatin regulatory proteins.

The synergistic effect of these two drugs is dependent on NQO1 expression, further supporting the cancer cell selectivity of this dual agent treatment. Tumor-specific targeting by IB-DNQ and Rucaparib combination treatment provides strong support for their potential use as a clinical therapy for patients with TNBC and our findings provide a compendium of cellular signaling changes with short single-agent and combination treatments defining the pathways that are highly affected that could contribute to cancer resistance mechanisms.

## RESULTS

### Rucaparib sensitizes TNBC cells to IB-DNQ-induced cell death in an NQO1 dependent manner

Previous studies determined that dual treatment of lung, pancreas, and breast cancer cell types with β-lapachone and Rucaparib leads to increased cancer cell death compared to either treatment alone, indicating synergistic lethality (12). However, since IB-DNQ has demonstrated greater potency than β-lapachone in some cancer cell lines, we wanted to perform a dose-dependent analysis of TNBC survival following IB-DNQ single agent or Rucaparib combination treatment for 4 hours. The mechanisms that underlie IB-DNQ and Rucaparib single agent and combination treatment on system-wide protein abundance and function and the downstream mechanisms that lead to cell death are needed to understand differences in cellular response to these therapies. To achieve this, we performed global proteomics using the MDA-MB-231 TNBC cell lines after performance of dose-response studies to determine optimal treatment concentrations for use in our assessment of IB-DNQ mechanisms of action.

To determine optimal treatment conditions, TNBC cell survival was assessed 7 days after the 4-hour treatment as indicated by relative DNA content measured via Hoechst and fluorescent quantification. We analyzed cell survival of three triple negative cell lines, MDA-MB-231, HCC1937, and MDA-MB-468, with increasing concentrations of IB-DNQ or pretreatment with nonlethal concentration of 15 µM Rucaparib as previously shown (12). Many TNBC cell lines have been shown to overexpress NQO1 including HCC1937 cells (11). For HCC1937 cells, we also performed combination treatment with 50 µM Dicoumarol to inhibit NQO1 activity (28,29). As shown in Figure 1c, HCC1937 cells were sensitive to nM concentrations of IB-DNQ. Pretreatment inhibition of NQO1 with Dicoumarol greatly reduced the effectiveness of IB-DNQ, illustrating strong NQO1-dependence for IB-DNQ cell killing (Figure 1c). Combination treatment of 15 µM Rucaparib along with IB-DNQ showed effective HCC1937 cell killing at concentrations around 100-200 nM. We extended these studies to other TNBC cell lines and used an exogenous expression approach to control the cellular levels of NQO1. Both the MDA-MB-231 and MDA-MB-468 cell lines contain a polymorphism in NQO1 that lead to its rapid degradation (NQO1*2) (30). The presence of this NQO1 variant allows us to perform isogenic control studies on NQO1 dependence through use of NQO1 overexpression in these cells by a plasmid (NQO1+ cells). NQO1*2 TNBC cells showed strong resistance to IB-DNQ treatment alone compared to their isogenic counterpart where NQO1 was overexpressed (Figure 1d and Supplementary Figure 1).

Furthermore, addition of non-lethal concentration of Rucaparib (15 µM) in MDA-MB-231 and MDA-MB-468 TNBC cell lines greatly potentiated the therapeutic effect of IB-DNQ. Cell death occurred in an NQO1-dependent manner at lower relative IB-DNQ concentrations than needed for cell killing with IB-DNQ alone in combination treatment with Rucaparib in all TNBC cell lines tested (Figure 1c-d, Supplementary Figure 1). Prior studies on MDA-MB-231 cells required combination treatment of 3 µM β-lapachone with 15 µM Rucaparib for cell killing whereas 0.1 µM IB-DNQ was sufficient for TNBC cell killing in combination with 15 µM Rucaparib in our long-term survival assays (12).

### Characterizing global protein and phosphorylation dynamics following IB-DNQ and Rucaparib treatment

The mechanisms that underlie IB-DNQ and Rucaparib single agent and combination treatment on system-wide protein abundance and function and the downstream mechanisms that lead to cell death are needed to understand differences in cellular response to these therapies. To achieve this, we performed global proteomics using the MDA-MB-231 TNBC cell lines that contain a polymorphism in NQO1 that lead to its rapid degradation (NQO1*2), and isogenic NQO1+ cells where wild-type NQO1 is stably overexpressed. Four biological replicates of each cell line were grown and treated with a sublethal low dose (0.1 µM) of IB-DNQ, a lethal high dose (0.4 µM) of IB-DNQ, 15 µM of Rucaparib (non-lethal concentration), or a combination of low dose IB-DNQ and Rucaparib for 4 hours. Cells were isolated for proteomics analysis following the four-hour treatment to facilitate assessment of rapid signaling changes to IB-DNQ treatment without confounding variables associated with cell death at later time points. Following the specified treatments, cells were collected and lysed and resulting global protein abundance was analyzed using LC-MS/MS. A total of 8,070 proteins were identified across all treatment groups and genotypes. Overall, only 18 proteins had significant changes (p ≤ 0.05) in abundance with the 4-hour drug treatments compared to the control in NQO1+ cells; no proteins changed in NQO1*2 cells reinforcing the selectivity of the NQO1-bioactivatable agent IB-DNQ (Supplementary Figure 2). The low amount of protein abundance changes following drug treatment suggested that other factors such as alterations in protein modifications and / or function would be more descriptive of the rapid TNBC changes associated with decreased long-term cell survival in the various treatment conditions.

Phosphorylation of proteins is one of the most abundant and well characterized protein post-translational modifications that occurs rapidly and is likely to be altered by IB-DNQ and/or Rucaparib 4-hour drug treatment. Since we observed a synergistic effect of TNBC treatment with both low dose IB-DNQ and Rucaparib, we expect DNA damage to occur as previously observed for equitoxic 3 µM β-lapachone treatment (12,13).

Phosphorylation of proteins is a major contributor to the signaling and recruitment of DNA repair proteins (31,32). Therefore, we performed phosphoproteomics to determine if there are changes in the phosphorylation state of proteins involved in DNA repair and / or other cellular pathways. Phosphoproteomics using MS^3^-based tandem mass tag (TMT) reporter ion quantitation identified a total of 5,863 proteins and 17,102 phosphopeptide groups (31,372 total peptide groups) across all NQO1+ treatment groups in our phosphoproteomics dataset (Figure 2 and Supplementary Figure 3) and 4,371 proteins and 12,206 phosphopeptide groups (18,486 total peptide groups) across all treatment groups in NQO1*2 cells (Supplementary Figure 4). Principle component analysis of NQO1+ samples shows that the DMSO and 15 µM Rucaparib biological replicate studies clearly group together, while the sublethal 0.1 µM IB-DNQ, lethal dose 0.4 µM IB-DNQ, and combination treatment replicates separate away from these groups (Supplementary Figure 3a). Rucaparib grouping with DMSO is not surprising because, without induction of DNA damage, we do not expect much of an effect from the drug obstructing DNA repair in MDA-MB-231 TNBC cells, which do not harbor BRCA mutations (33). This is also consistent with Figure 1d which shows that 15 µM Rucaparib single agent treatment is non-lethal (12). Similarly, there is only a small separation between the replicate groups for DMSO and the sublethal IB-DNQ dose alone. Interestingly, the phosphoproteomics data for 0.4 µM IB-DNQ and combination treatments group distantly from one another, consistent with the hypothesis that these two drug treatments lead to cell death through distinct molecular mechanisms.

**Figure 2:**
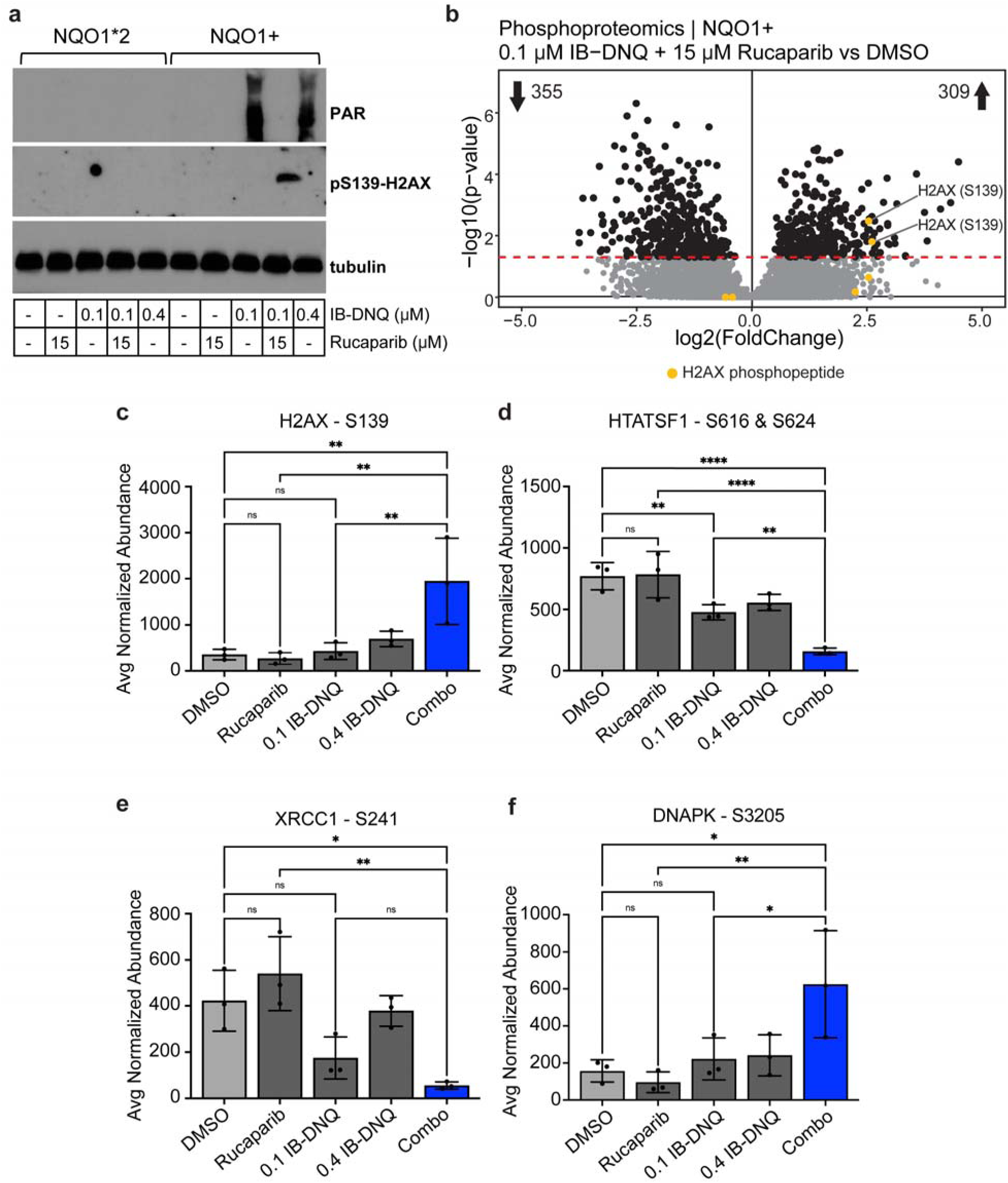
DNA damage response is increased in combination treated cells compared single agent and DMSO treatments. A) Western blot analysis of PAR and phospho-serine 139 of H2AX in NQO1*2 and NQO1+ cells. Tubulin was used as the loading control. IB-DNQ and Rucaparib treatments for each lane are shown in the table. b) Volcano plot shows differential phosphorylation of phosphopeptide groups in 0.1 µM IB-DNQ + 15 µM Rucaparib treated NQO1+ cells compared to DMSO treated cells. Yellow points represent phosphopeptides of histone protein H2AX. Red dashed line is p-value cut off p=0.05. c-f) Bar graphs of average normalized phosphopeptide abundance associated with DNA damage response proteins H2AX (c), HTATSF1 (d), XRCC1 (e), DNAPK (f). Phosphorylated residue is shown above each graph. Light gray bars show abundance in DMSO, dark gray bars show abundance in single agent treatments, and blue bars show abundance for the IB-DNQ and Rucaparib combination treatment. Comparisons between groups show statistical significance as calculated by one-way ANOVA and Tukey multiple comparisons test. ns = not significant, * = p<0.05, ** = p<0.01, *** = p<0.001, **** = p<0.0001.

Throughout NQO1*2 samples where NQO1 is rapidly degraded, we observe a similar number of phosphorylation changes in each treatment group with the greatest number of significant changes resulting from combination treatment (25 phosphopeptides increasing and 28 decreasing in abundance, Supplementary Figure 4). In NQO1+ cells, we observe more distinct phosphorylation changes in each of the different IB-DNQ treatments (Figure 2b and Supplementary Figure 3c-e). Specifically, the low dose IB-DNQ and Rucaparib combination treatment leads to the greatest number of significant phosphopeptide changes with 309 increasing and 355 decreasing (Figure 2b). Gene ontology analysis of the proteins with significantly changing phosphorylation shows significant enrichment of pathways involved in DNA repair, RNA processing, and other processes consistent with DNA damage response, cell cycle disruption, and alterations in RNA transcription pathways (Supplementary Figure 3b). The larger number of phosphorylation changes observed with NQO1+ compared to NQO1*2 shows that MDA-MB-231 TNBC cells required NQO1 for broad impacts on phosphorylation by IB-DNQ + Rucaparib combination treatment. This result supports the characterization of IB-DNQ as an NQO1 bioactivated drug. We also observe a synergistic effect of combining the sublethal 0.1 µM IB-DNQ and 15 µM Rucaparib treatments which supports the potential use of these drugs in an effective combination therapy for tumors with high expression of NQO1.

### Dual agent treatment induces DNA damage response

Histone protein H2AX phosphorylation at serine 139 (γH2AX) is a canonical marker for DNA damage (reviewed in (34,35)). The Ser 139 residue of H2AX is phosphorylated by the kinases ATM and ATR (36,37). By western blot, we observe increased pSer 139-H2AX with the IB-DNQ + Rucaparib combination treatment in an NQO1-dependent manner indicating DNA damage in these cells (Figure 2a). Although pS139-H2AX was not observed in single agent IB-DNQ treatments at 0.1 and 0.4 µM IB-DNQ, PARP hyperactivation did occur as shown by detection of protein PARylation (i.e. poly(ADP-ribosyl)ation) in a fully NQO1-dependent manner. PARP hyperactivation was not observed in co-treatment of 0.1 uM IB-DNQ with Rucaparib because of PARP inhibition (Figure 2a). Our phosphoproteomics analysis readily confirmed significant increases in the phosphorylation of H2AX at S139 is in NQO1+ cells treated with the combination drug but not with any of the single agent treatments (Figure 2b-c and Supplementary Figure 3c-e). Additionally, 2-way ANOVA analysis of 0.1 µM Rucaparib and IB-DNQ H2AX modification levels relative to combination treatment revealed a clear synergistic increase to combination treatment (Supplementary Table 15, interaction p = 0.0209).

Phosphoproteomics analysis of combination treatment induced changes allows for additional insights into pathway changes associated with H2AX phosphorylation increases. Additionally significant phosphorylation changes were observed in DNA damage response proteins HTATSF1, XRCC1, and DNAPK in TNBC cells treated with the combination compared to DMSO (Figure 2d-f). HTATSF1 is a component of the 17S US SnRNP complex of the spliceosome and plays a role in double-strand break repair in a phosphorylation-dependent manner (38–41). HTATSF1 also interacts with PARP modified poly-ADP-ribosylated RPA1 (41). Phosphorylation of HTATSF1 at Ser residues 616 and 624 were significantly decreased by single agent treatment with IB-DNQ at both the low and high dose treatment levels relative to DMSO (Figure 2d). Additionally, this HTATSF1 phosphorylation event showed a synergistic decrease in response to combination treatment with of 0.1 µM IB-DNQ and 15 µM Rucaparib for 4-hours through unknown mechanisms (Supplementary Table 15, interaction p=0.0344). XRCC1 acts as a scaffold to mediate assembly of DNA repair complexes and negatively regulates PARP1 activity (42–44). XRCC1 phosphorylation at Ser 241 was significantly decreased by 0.1 µM IB-DNQ single agent treatment (Figure 2e). Higher dose 0.4 µM IB-DNQ treatment did not significantly change XRCC1 Ser 241 levels, suggesting a differential signaling response although PARP hyperactivation occurs in both treatment conditions (Figure 2a). Inhibition of PARP in the combination treatment did not further alter XRCC1 Ser 241 levels relative to 0.1 µM IB-DNQ treatment (Figure 2e). DNAPK is a kinase that senses DNA damage and is required for non-homologous end joining DNA repair (45–49). Like H2AX phosphorylation at Ser 139, DNAPK phosphorylation at Ser 3205 showed significant synergistic increases in combination treatment (Figure 2f, Supplementary Table 15, interaction p=0.0369). Phosphorylation changes across these important DNA repair proteins is indicative of signaling changes related to DNA damage and repair response, however, many of these site-specific changes have not been previously characterized and could be used to delineate specific pathway responses to single and combination treatments with IB-DNQ.

### Kinase substrate enrichment analysis predicts changes in DNA damage response, cell cycle, and transcriptional regulation

To determine the kinases associated with the changing phosphorylation sites, we performed computational analysis using Kinase Substrate Enrichment Analysis (KSEA) (50). KSEA predicts kinase activity from phosphoproteomics datasets by utilizing prior knowledge of kinase-substrate relationships from curated databases and computational prediction tools (50–53). Each kinase is scored based on the relative hyper- and hypo-phosphorylation of its substrates, as identified from PhosphoSitePlus and NetworkIn (52,54,55). The positive and negative values represent an increase or decrease in inferred kinase activity in response to drug treatment relative to DMSO control, and blue and red bars in the bar plots represent kinases with significantly changing activity (p ≤ 0.05, Figure 3a and Supplementary Figure 5). With 0.1 µM IB-DNQ treatment, PRKCB was the only kinase with decreased activity, while 12 kinases showed increased activity (Supplementary Figure 5a). Single-agent Rucaparib reduced the activity of PRKCB and CDK7 (Supplementary Figure 5b). At 0.4 µM IB-DNQ, four kinases increased and two decreased in activity (Supplementary Figure 5c). With combination treatment, more kinase activity is impacted with 3 kinases have significantly decreased activity and 14 kinases have increased activity (Figure 3a). CDK1 and CDK2, which are essential for promoting cell cycle progression (reviewed in (56)) are predicted to have significantly decreased activity. Decreased activity of CDK1 and CDK2 is consistent with stalling of the cell cycle, which is known to occur during the DNA damage response (57). NEK2, AURKC, RPS6KA2, PAK2, and PLK1 are also involved in cell cycle regulation reviewed in (58–62) and are predicted to have significantly increased kinase activity. ATM and CHEK1, which positively regulate the DNA damage response (56,63), also have significantly increased activity, which aligns with increased DNA damage we expect from the drug treatment and phosphorylation site changes such as H2AX Ser 139.

**Figure 3:**
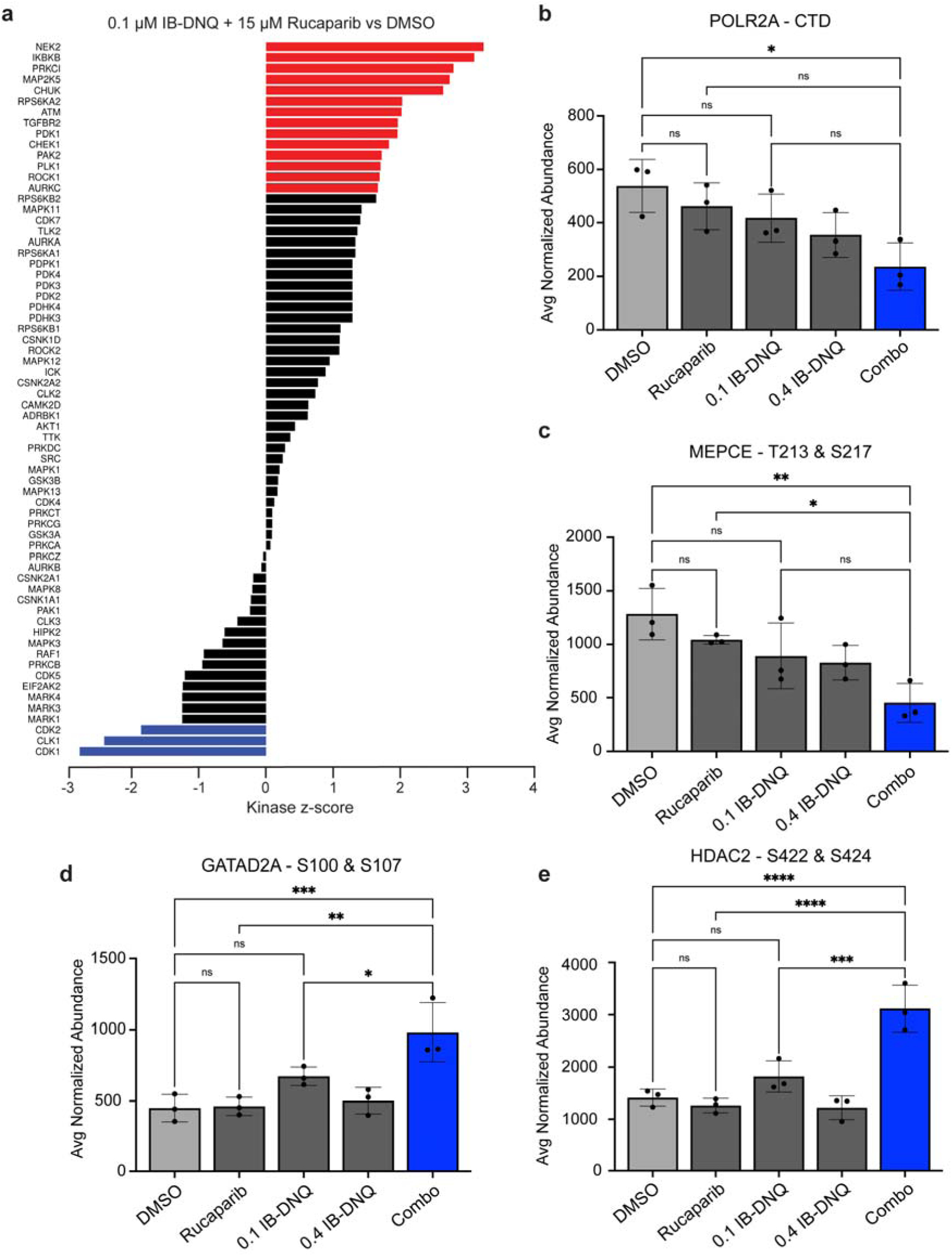
Bioinformatic analysis of phosphoproteomics reveals changes in DNA damage response, cell cycle, and transcriptional regulation. a) Kinase substrate enrichment analysis (KSEA) of combination drug treated cells compared to DMSO. KSEA bar plot for 0.1 µM IB-DNQ + 15 µM Rucaparib. Red bars indicate kinases with significantly increased inferred activity (p ≤ 0.05), and blue bars indicate kinases with significantly decreased inferred activity. Black bars show kinases with differential inferred kinase activity but are not statistically significant. b-e) Bar graphs of average normalized phosphopeptide abundance associated with transcription regulatory proteins POLR2A (b), MEPCE (c), GATAD2 (d), HDAC2 (e). Phosphorylated residue is shown above each graph. The sequence detected for the C-terminal domain (CTD) of POLR2A is YSPTSPKYSPTSPK where the serines are differentially phosphorylated. Light gray bars show abundance in DMSO, dark gray bars show abundance in single agent treatments, and blue bars show abundance for the IB-DNQ and Rucaparib combination treatment. Comparisons between groups show statistical significance as calculated by one-way ANOVA and Tukey multiple comparisons test. ns = not significant, * = p<0.05, ** = p<0.01, *** = p<0.001, **** = p<0.0001.

Together these findings suggest that IB-DNQ-induced DNA damage is being repaired in the absence of Rucaparib treatment, and the combination drug treatment leads to an accumulation of DNA damage.

Our findings from KSEA analysis motivated the identification of additional phosphorylation targets with significant changes in single agent and/or combination treatments in our study. The target amino acid sequence for all cyclin dependent kinases (CDKs) is identical, which includes S/TPXK/R, where S or T is phosphorylated and X is any amino acid (64–68). Several CDKs are identified through KSEA indicating potential alteration of the activity of any kinases within the CDK family. An important target of CDK activity is the RNA transcription machinery, including the key regulator of mRNA and long noncoding (lnc)RNA production, RNA polymerase II. In our single and combination treatment studies, we identified alterations in RNA Polymerase phosphorylation typically associated with CDK7 and CDK9 rather than CDK1/2.

Specifically, we detected a significant decrease in phosphorylation of the regulatory C-terminal domain (CTD) of RNA Polymerase II largest subunit POLR2A following combination drug treatment compared to DMSO (Figure 3b) CTD phosphorylation occurs within repetitive amino acid repeats in one of the catalytic core subunits of RNA polymerase II, and decreasing phosphorylation of the regulatory CTD has previously been shown to correlate with decreased transcriptional activity (69). Although not statistically significant, RNA Polymerase II CTD phosphorylation trends towards decreased with single agent treatment (Figure 3b). MEPCE is a 5’ methylphosphate capping enzyme which is part of the P-TEFb complex which also contains CDK9 (70). The P-TEFb complex phosphorylates the RNA Polymerase II CTD to facilitate transcription elongation. MEPCE has reduced phosphorylation at Thr 213 and Ser 217 following combination drug treatment (Figure 3c) and closely mirrors the changes observed in RNA Polymerase II CTD phosphorylation, possibly indicating the CDK9 containing P-TEFb complex in the changes observed in both transcription related targets (Figure 3b). Consistent with an overall decrease in RNA Polymerase II transcription during the DNA damage response, significant changes in phosphorylation of chromatin related complexes were also observed. These chromatin regulators include transcriptional repressors and components of the histone deacetylase complex NuRD: GATAD2A and HDAC2 (Figure 3d-e) (71–73). GATAD2A is required for PAR-dependent recruitment of the NuRD complex to DNA double-strand breaks (74).

GATAD2A phosphorylation at Ser 100 and 107 significantly increases in the combination treatment relative to DMSO or single agent treatment with 0.1 µM IB-DNQ or 15 µM Rucaparib. Histone deacetylase HDAC2 phosphorylation at Ser 422 and 424 also significantly increases only in combination treatment conditions. Together, these data suggest that RNA transcription and chromatin-related pathways are also directly altered specifically because of IB-DNQ + Rucaparib combination treatment in an NQO1-dependent manner.

### Protein thermal stability impacted by single and dual agent treatments

Proteome-wide changes to protein-protein interactions, post-translational modifications, folding, and other biophysical properties of proteins can be assessed using thermal proteome profiling (TPP) (75–79). TPP is used to determine thermal stability of each protein under different conditions, which are then analyzed computationally to identify proteins and mechanisms impacted by treatment (80,81). These data could be highly complementary to phosphoproteomics findings but could also provide us with novel mechanistic insights since neither method has previously been performed in cancer models with treatment using NQO1 bioactivatable therapeutics. We analyzed our TPP data using the R-package InflectSSP to fit data to melt curves, calculate melt temperatures (T_m_), and identify statistically significant shifts in protein melt temperature (81). TPP has been widely used to determine drug-target engagement (79) and changes in protein-protein interactions (75,82), therefore we can apply it to test and compare the proteome-wide impacts on protein thermal stability across individual drug treatments as defined by significant melt shifts. Compared to DMSO control, 713 proteins significantly changed in stability when treated with 0.1 µM IB-DNQ, 1541 proteins changed in stability when treated with 0.4 µM IB-DNQ, 1824 proteins changed in stability with 15 µM Rucaparib, and 135 proteins significantly changed in thermal stability with combination treatment (Supplementary Figure 6a). With Rucaparib treatment, PARP1 appears to trend towards decreased thermal stability compared to DMSO treatment, but this was not statistically significant (p=0.0646; Supplementary Figure 6b). NQO1 thermal stability is unchanged when cells are treated with 0.1 or 0.4 µM IB-DNQ (Supplementary Figure 6c-d). This is likely due to the fact that IB-DNQ is a futile substrate for NQO1, and the transient IB-DNQ-NQO1 interaction is not sufficient to alter average NQO1 thermal stability because of a rapid engagement time.

### Combination drug treatment increases H2AX phosphorylation and decreases stability

InflectSSP analysis identified significant thermal stability changes in the histone protein H2AX, which was significantly altered in specific treatment regimens of IB-DNQ (Figure 4). The average melt temperature for H2AX in DMSO was determined to be 64.9°C across all our experiments (Figure 4). Significant destabilization of H2AX was observed in TNBC cells with high dose 0.4 µM IB-DNQ with a decrease in melt temperature of 8.53°C as calculated by InflectSSP (Figure 4b, p=0.0465). We also observed significant destabilization of H2AX with the combination treatment where the melt temperature decreased similarly by 8.75°C (Figure 4d, p=0.0193). In prior work, PARP has been shown to bind to nucleosomes and to have preferential binding to nucleosomes that contain H2AX relative to H2A by surface plasmon resonance (83,84). However, treatment with the PARP inhibitor Rucaparib in single agent treatment was not sufficient to significantly alter H2AX thermal stability even though the overall change in melt temperature trended towards destabilized (Figure 4c). H2AX is readily phosphorylated as shown in Figure 2, but only in combination treatment conditions in the TNBC cells at the 4-hour treatment time point, so phosphorylation-induced H2AX protein biophysical changes alone do not explain these findings. However, the two treatment conditions in which H2AX thermal stability was substantially altered in MDA-MD-231 cells are treatment conditions associated with poor TNBC cell survival 7 days after the 4-hour treatment as indicated (Figure 1d). Overall, these results suggests that lethal IB-DNQ treatment conditions as a single agent or in combination with Rucaparib inhibition significantly alter H2AX thermal stability. It is likely that these alterations occur as a consequence of changes in the DNA repair and/or chromatin-related protein machinery and their associations with H2AX induced by NQO1 bioactivated compounds. This could be due to precise biophysical properties of H2AX or due to changes in protein-protein interactions, protein modifications, and / or overall cell metabolic state that occur following IB-DNQ treatment (10). These data in combination with our cell survival studies (Figure 1) also suggest that significant changes in H2AX thermal stability could be an early indicator of cell sensitivity to NQO1-bioactivated compound therapies.

**Figure 4:**
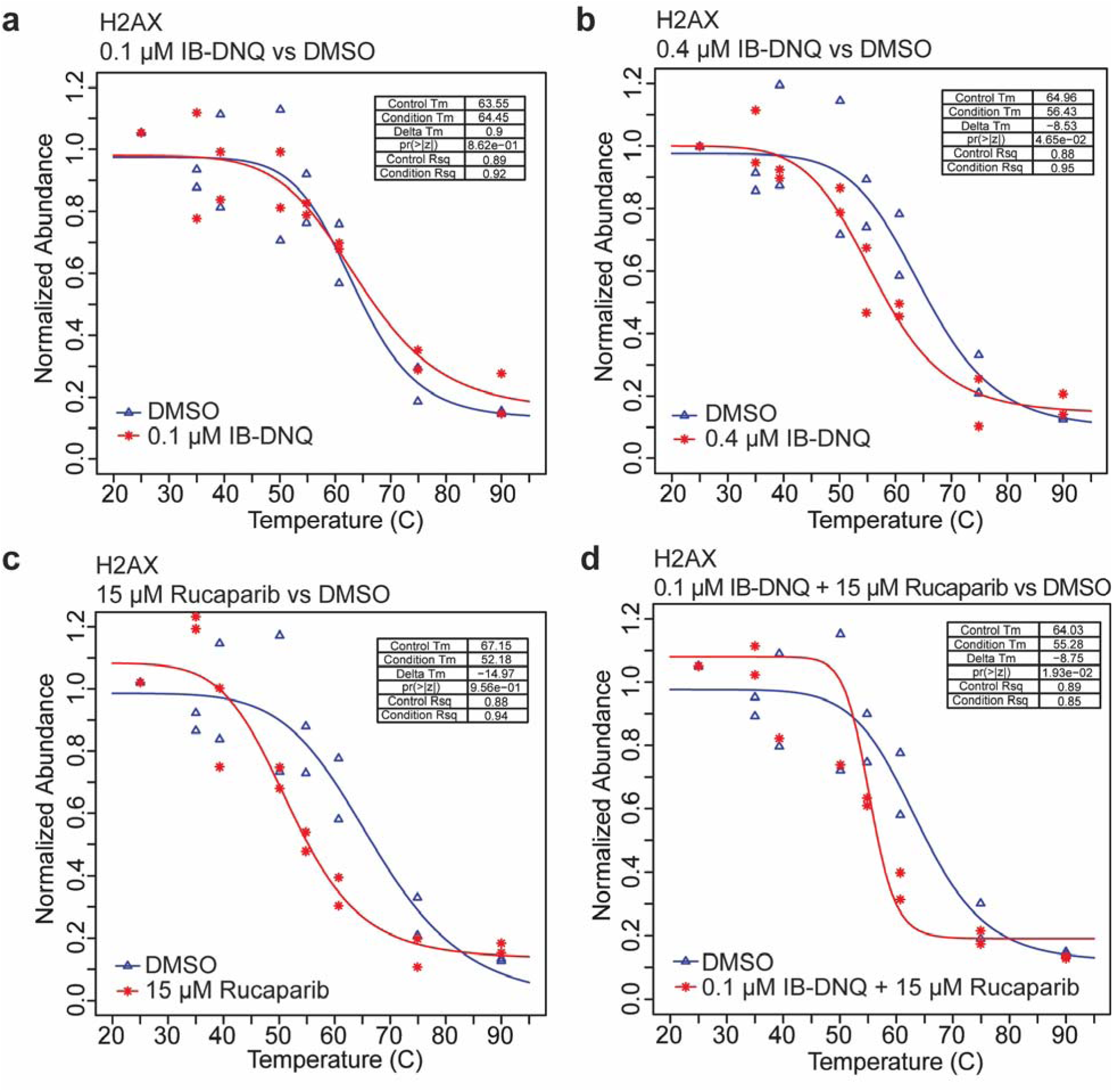
Thermal stability of H2AX following different drug treatments. a-d) Melt curves for H2AX in 0.1 µM IB-DNQ (a), 0.4 µM IB-DNQ (b), 15 µM Rucaparib (c), and combination drug (d) treated cells compared to DMSO as calculated by InflectSSP. Blue lines represent melt curves for H2AX in DMSO treated cells, and red lines represent melt curves for H2AX in drug treated cells. Melt temperature (Tm) is shown for control (DMSO) and condition (drug treatment). Difference in melt temperature (delta Tm), calculated p-values, and R^2^ of curve fit are also shown.

### DNA repair and associated proteins have increased thermal stability following combination treatment

TPP data has utility for identification of protein changes in thermal stability as performed using melt-shift analysis tools like InflectSSP, but studies focused on protein-protein interaction alterations have also greatly benefitted from analysis using thermal proximity coaggregation analysis (TPCA) (75,85). While melt shift analysis helps identify proteins with biophysical changes, TPCA is a method that more specifically monitors binary protein interactions and protein complex dynamics (75). For TPCA analysis of the IB-DNQ and Rucaparib studies in TNBC cells, we employed the Rtpca R-package which uses annotated protein interaction databases (CORUM and BioGRID were used here) and measures Euclidean distance between protein melt curves to identify protein-protein interaction changes (86). TPCA of combination drug treatment compared to DMSO TPP data reveals changes in protein-protein interactions and protein complex assembly that are described by TPCA coaggregation scores for all binary protein interactions tested relative to measures of their statistical significance (-log_10_(p-value)) in Figures 5a-b. Coaggregation score is determined by comparing the average melt curve distance in drug and DMSO (i.e. distance in drug – distance in DMSO). Therefore, positive coaggregation scores indicate decreased protein binary interaction while negative coaggregation scores indicate an increase in binary interactions. Proteins involved in DNA repair processes are highlighted in blue (Figure 5a). Upon in-depth review of the TPCA results, we identified multiple subunits of the SWI / SNF complex, a protein complex involved in chromatin remodeling during DNA replication, transcription, and DNA repair (87), were significantly changing in their binary interaction strength within the complex (Figure 5b, highlighted in orange). We were specifically interested in the subset of SWI / SNF proteins that were increasing in interactions with one another as this could indicate tighter association of protein complexes and proteins that work together resulting in enhanced function. In DMSO, curves for these proteins are loosely associated, but in combination drug treatment, curves become more tightly associated indicating a stronger interaction of the SWI / SNF subunits in the combination drug treatment (Figure 5c). Additional comparison of these findings to the phosphoproteomics datasets identified an increase in phosphorylation of SMARCC2 at Thr 548 following combination drug treatment (Figure 5d). Although not identified in the group of SWI / SNF subunits decreasing in stability, phosphorylation of SMARCAD1 at Ser 214 is decreasing following combination drug treatment (Figure 5e). Our TPCA findings suggest that SWI / SNF complex associations are enhanced following the combination drug treatment which could suggest changes in chromatin remodeling associated with extensive DNA damage.

**Figure 5:**
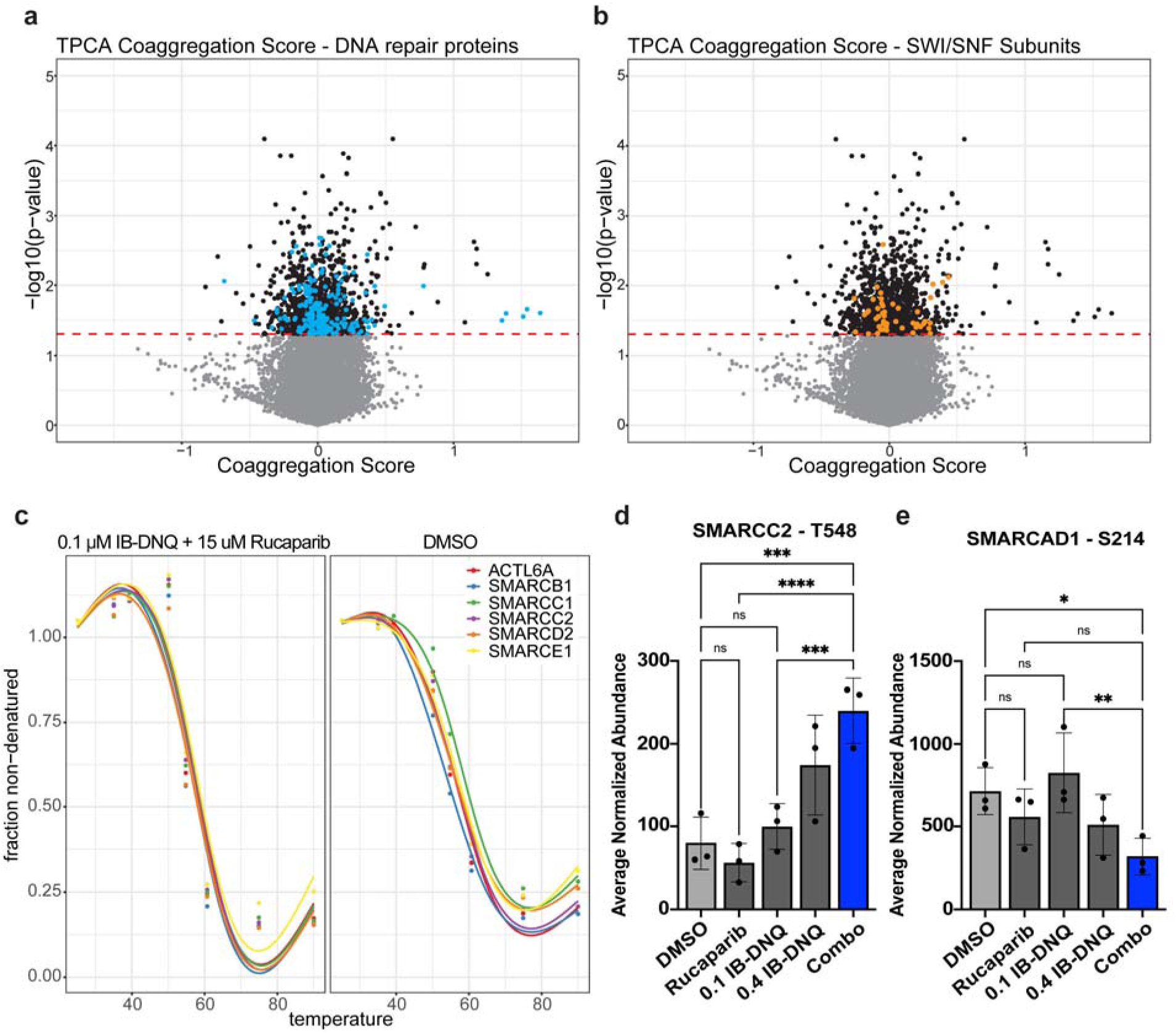
Thermal stability of DNA damage repair proteins and chromatin remodeling complex SWI / SNF are affected by combination drug treatment. a,b) Volcano plots of differential protein coaggregation as identified by thermal proximity coaggregation (TPCA) in combination treatment compared to DMSO. Blue dots indicate protein pairs with significantly changed coaggregation involved in DNA damage repair (a) and orange dots indicate proteins in the SWI / SNF complex with significantly changed coaggregation (b). Red dashed line is p-value cut off p=0.05. c) Melt curves for proteins involved in the SWI / SNF complex in DMSO and IB-DNQ + Rucaparib treated cells. Different colors represent the different SWI / SNF subunits. d,e) Average normalized abundance of phosphopeptides for SMARCC2 (d) and SMARCAD1 (e) in each of the treatment groups. Phosphorylated residue is shown above each graph. Light gray bars show abundance in DMSO, dark gray bars show abundance in single agent treatments, and blue bars show abundance for the IB-DNQ and Rucaparib combination treatment. Comparisons between groups show statistical significance as calculated by one-way ANOVA and Fisher LSD multiple comparison test. ns = not significant, * = p<0.05, ** = p<0.01, *** = p<0.001, **** = p<0.0001.

### Protein-protein interaction changes between cell cycle regulators

Our observations of differential kinase activity for several cell cycle regulatory proteins lead us to hypothesize that interactions between cell cycle regulators could be significantly altered in our IB-DNQ treatment conditions. Cell cycle has been well-described to have a heavy reliance on phosphorylation-based signaling for regulation of complex protein-protein interactions and protein turnover, which could also influence protein interaction status (88–90). Here, we highlight changes in TPCA with the central cell cycle kinase CDK1 (Figure 6). Figure 6a highlights binary protein interaction changes between proteins involved in cell cycle regulation (yellow). p53 and tubulin alpha-1A chain (TUBA1A), which interact at the mitotic centrosome (91,92), are shown to have decreased association by TPCA with the combination drug treatment suggesting decreased interaction (Figure 6a and c, p = 0.0066). Several binary protein interactions between CDK1 targets are changing in TPCA calculated distance, suggesting wide-spread changes in CDK1 function following IB-DNQ and Rucaparib combination treatment (Figure 6b, green points). Nine binary interactions between CDC20 and known interactors have stronger interactions in the combination drug treatment compared to DMSO, indicating a potentially important signaling role for CDC20 in response to combination treatment. CDC20 is a key regulator of the ubiquitin ligase activity of the anaphase promoting complex that provides substrate specificity during metaphase/anaphase (93). Protein-protein interaction network analysis using STRING provides visual representation of the relevant pathways (i.e. Gene Ontology biological processes) these protein interactions are significantly enriched in with CDC20 during combination treatment conditions specifically (Figure 6d). These proteins are involved in chromosome organization, regulation of mitosis, and microtubule organization. These findings reveal specific protein-protein interaction changes that occur during IB-DNQ / Rucaparib combination treatment in TNBC cells likely associated with key cell cycle alterations. These types of mechanistic details can provide additional insights into TNBC cell vulnerabilities that could be targeted through the development of other IB-DNQ-based combination therapies.

**Figure 6:**
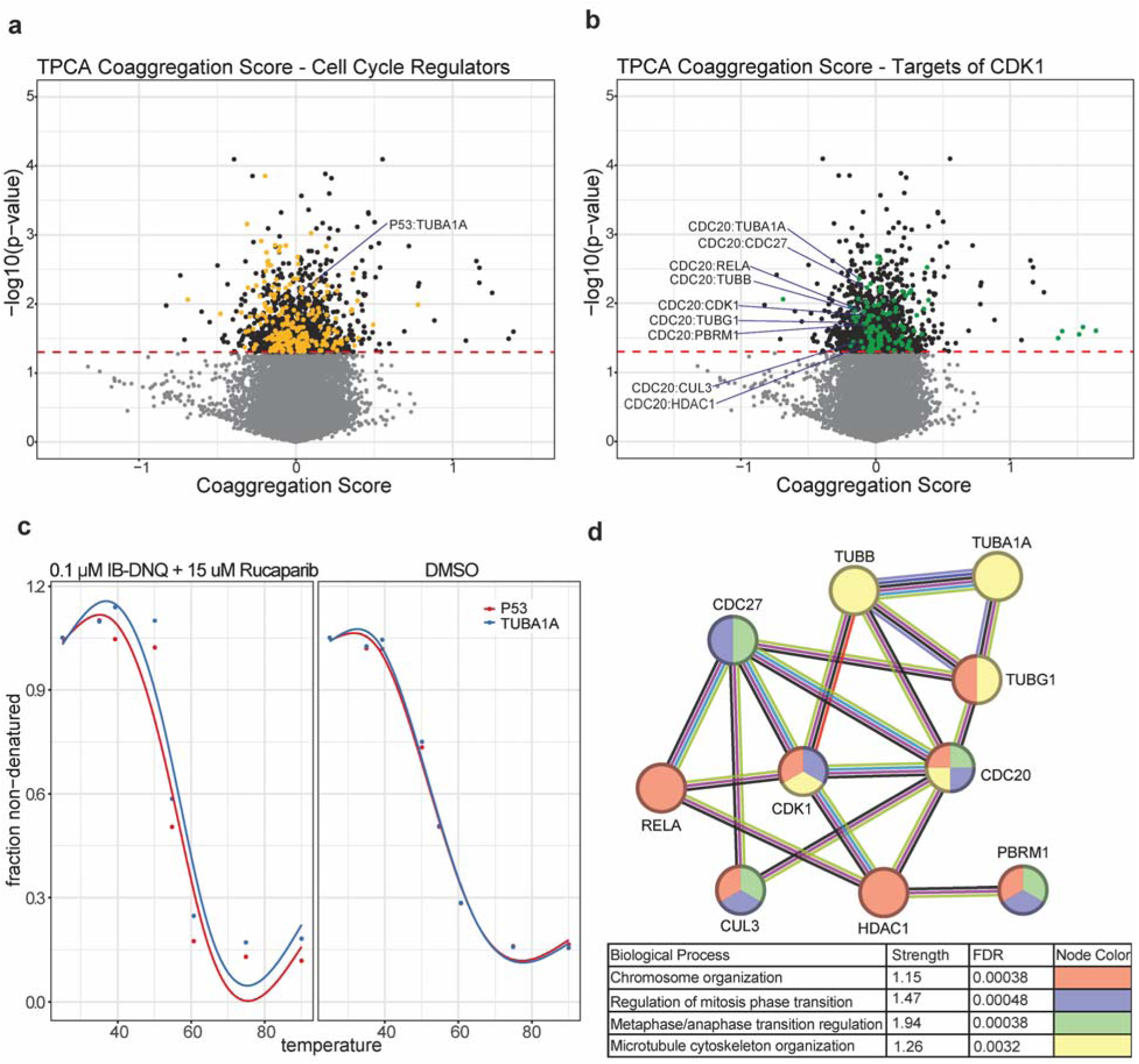
Thermal stability of cell cycle regulators and CDK1 targets are affected by combination drug treatment. a,b) Volcano plots of differential protein coaggregation as identified by thermal proximity coaggregation (TPCA) in combination treatment compared to DMSO. Yellow dots indicate protein pairs with significantly changed coaggregation involved in cell cycle regulation (a) and green dots indicate targets of CDK1 with significantly changed coaggregation (b). Protein pairs labeled in (b) include CDC20 as an interaction partner (TUBA1A p=0.0044; CDC27 p=0.0059; RELA p=0.0132; TUBB p=0.0129; CDK1 p=0.0152; TUBG1 p=0.0182; PBRM1 p=0.01945; CUL3 p=0.0358; HDAC1 p=0.0440). Red dashed line is p-value cut off p=0.05. c) Melt curves for p53 (red line) and TUBA1A (blue line) in DMSO and combination treated cells (p=0.0406). d) Protein-protein interaction network of proteins that are changing in interaction with CDC20. String analysis shows these proteins are involved in chromosome organization, regulation of mitosis, and cytoskeleton organization, all of which can be related back to cell cycle. Node color corresponds to the biological process the protein is involved in; color of lines connecting nodes corresponds to origin of protein-protein interaction annotation (black=co-expression, pink=experimentally determined, light blue=from curated database, green=text mining, purple=protein homology, red=gene fusions, dark blue=gene co-occurrence).

### Dual and single treatments have distinct mechanisms leading to cell death

Single agent treatment with high dose IB-DNQ and combination treatment with Rucaparib are both cancer cell death-inducing treatments for multiple TNBC cell lines expressing high levels of NQO1 (Figure 1 and Supplementary Figure 1). In our phosphoproteomics and TPP analyses, we have identified several pathways which were altered specifically in treatment conditions which resulted in cell death. To comprehensively annotate changes that were specifically identified in IB-DNQ treatment conditions and in TNBC cell-killing conditions, we performed a direct multiomics comparison between the 0.4 µM IB-DNQ and 0.1 µM IB-DNQ / 15 µM Rucaparib combination treatment to classify both shared and unique significant changes associated with the early 4-hour response to each lethal IB-DNQ treatment condition. Figure 7a compares significant changes in protein thermal stability between the two lethal treatment groups at the four-hour time point. A total of 22 overlapping proteins were identified with 16 proteins being significantly destabilized in the high dose IB-DNQ treatment that are stabilized in the combination treatment. These data support the hypothesis that high-dose NQO1-bioactivatable treatment occurs through distinct mechanisms relative to low dose IB-DNQ combination treatment with Rucaparib (Figure 1A) (12). For clarity, all significant changes were included with no consideration of the direction of change, however; all directionality information is available (Supplementary Tables 1-2). Changes in phosphorylation of peptides between the two lethal treatment groups of 0.4 µM IB-DNQ and 0.1 µM IB-DNQ / 15 µM Rucaparib were also compared (Figure 7b). There are 34 peptides that have significantly decreased phosphorylation and 16 peptides with significantly increased phosphorylation shared in both treatment groups (Figure 7b). Several of the proteins with phosphopeptides that are increasing with IB-DNQ/Rucaparib combination treatment and decreasing with 0.4 µM IB-DNQ are involved in RNA splicing regulation, linking specific changes in the RNA Polymerase II transcription process with different IB-DNQ treatment regimens. Proteins with significantly altered phosphorylation sites identified in the multiomics analysis include DDX23, SRSF7, SRRM2, SRRM1, and CDK12. CDK12 is a known direct regulator of phosphorylation at the RNA Polymerase II C-terminal domain which also significantly changes in combination IB-DNQ / Rucaparib treatment (Figure 3b). CDK12 phosphopeptide levels with modifications at Serine residues 332, 333, and 334 significantly decrease in the 0.4 µM IB-DNQ treatment while significantly increasing with 0.1 µM IB-DNQ + 15 µM Rucaparib treatment (Supplementary Figure 7b). CDK12 is also known to be essential for DNA repair through homologous recombination (HR) and was shown to suppress intronic polyadenylation events in key DNA repair factor genes (94). Our findings that CDK12 phosphorylation at Serine residues 332, 333, and 334 are differentially regulated in IB-DNQ treatment conditions suggests that CDK12 phosphorylation could also contribute to its role in the regulation of HR gene expression. Phosphorylation at two of the three CDK12 Serine residues 332, 333, and 334 was also significantly increased in 0.1 µM IB-DNQ + 15 µM Rucaparib treatment (Supplementary Figure 7c). Altogether, the overall low number of overlapping changes between the lethal treatments in TNBC cells again suggests that these regimens have distinct cellular consequences and interaction with the DNA damage response that led to cell death. Novel changes were observed for these pathways with uncharacterized effects on RNA Polymerase II transcription related processes.

**Figure 7:**
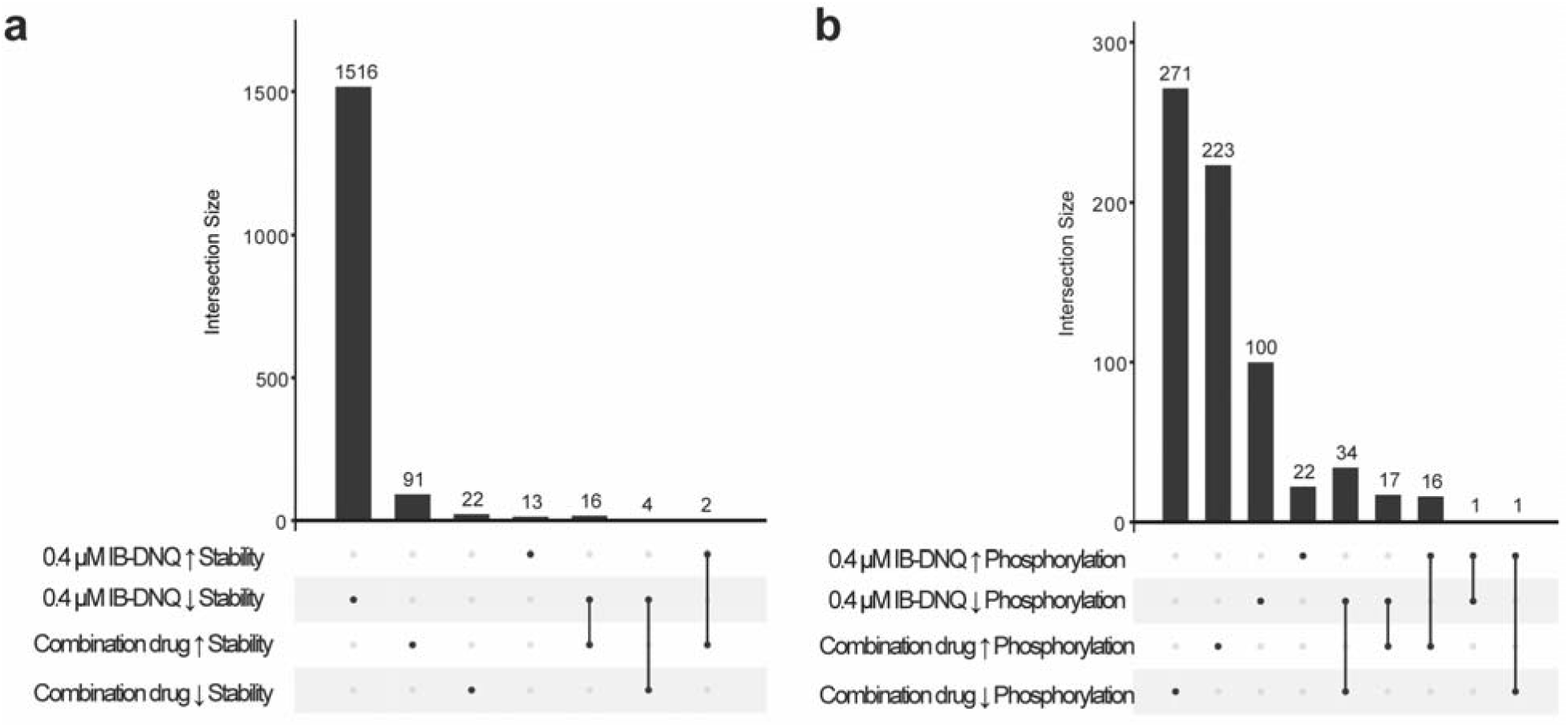
Comparison of protein stability changes and phosphorylation changes caused by lethal drug treatments 0.4 µM IB-DNQ or 0.1 µM IB-DNQ + 15 µM Rucaparib. a) Upset plot showing overlap between increased and decreased stability of proteins in the lethal drug treatment groups. b) Upset plot showing overlap of increased and decreased phosphorylation of peptides in the lethal drug treatment groups Intersection size is the number of unique or overlapping proteins or peptides in a group. Single dot represents proteins or peptides that are unique to that category, and 2 dots connected with a line show where there are overlapping proteins or peptides.

Finally, we also performed multiomics analysis for proteins with thermal stability and phosphorylation changes between the different IB-DNQ treatment groups - low dose, high dose, and combination (Supplementary Figure 7a). The largest number of target overlaps occurs between low dose and high dose IB-DNQ protein stability changes, suggesting reproducibility of many IB-DNQ cellular effects across treatment conditions. Examples of the proteins in this category include HDAC1, SMARCA1, and transcriptional regulators RPAP1, POLR1C, and CDK9. Interestingly, the overlap between changes in phosphorylation signaling and stability changes was limited across all treatment conditions suggesting that thermal stability analysis and phosphorylation provide unique molecular insights (77,95,96).

## DISCUSSION

TNBC accounts for a large portion of breast cancer diagnoses, and treatment for these patients is limited. Even with the most modern therapies, the risk of metastases and tumor recurrence is high (1,2). Research is ongoing to develop tumor-type specific treatment options that are highly effective with limited toxicity. In this work, IB-DNQ effects are highly NQO1-dependent in TNBC cells. We have carefully controlled for NQO1-dependence in our multiomics studies through use of TNBC cell lines in which we can control NQO1 expression. Cells with increased NQO1 activity do not survive high dose IB-DNQ treatment, and low dose IB-DNQ treatment substantially decreases TNBC cell survival with the addition of Rucaparib (Figure 1c-d and Supplementary Figure 1). Further, we observe greater number of changes to protein abundance and phosphorylation when NQO1 is increased compared to when NQO1 is low (Supplementary Figure 3-4 and Figure 2). These results strongly support the specificity of IB-DNQ for cells with increased NQO1 levels and strengthen the potential use of these drugs in TNBC patients for tumor selective therapy. The synergistic effect from the combination drug treatment is also supported by prior work studying TNBC treatment response to 3 µM β-lapachone and Rucaparib (12), although much lower concentrations of IB-DNQ are needed to achieve similar cellular effects, clearly demonstrating its increased effectiveness. Our highly quantitative multiomics studies also reveal numerous signaling and protein thermal stability changes that are altered through different drug response mechanisms with many, such as H2AX phosphorylation, showing a synergistic response to IB-DNQ + Rucaparib combination treatment (Supplementary Figure 3, and Figures 2, 3, & 5 Supplementary Table 15).

The predominant mechanisms found to be affected by the IB-DNQ + Rucaparib dual agent treatment are DNA damage response, cell cycle mechanisms, and RNA Polymerase II transcriptional regulation, which are highly targeted mechanisms in cancer treatments (97). The identification of precise sites of differential protein modification throughout our studies can serve as a resource for additional studies into the roles of various regulatory proteins on cellular events downstream of IB-DNQ treatment regimes.

Increased phosphorylation of H2AX at S139, which is indicative of increased DNA damage (34), has been observed previously upon treatment of NQO1+ cancer cells and NQO1+ xenograft tumor models with combination treatment of β-lapachone and Rucaparib (12). Our experiments replicate this finding with the IB-DNQ and Rucaparib combination treatment. We likely only observe this with the dual agent treatment because DNA damage is accumulating as PARP1 inhibition disrupts DNA damage repair. Additionally, the decreased stability of H2AX suggests alterations to chromatin organization, potentially for DNA repair or in preparation for apoptosis in response to the dual agent treatment. Our results indicate increased DNA damage and reduced DNA repair are key targets of the dual agent treatment, and together these suggest the cancer cells are unable to repair increased DNA damage leading to higher cell death, further supporting the potential use of this dual agent as TNBC therapeutic.

Analysis of protein-protein interactions using thermal profiling and TPCA revealed changes in interactions between proteins involved in the SWI / SNF complex following dual agent treatment. SWI / SNF is known to be involved in DNA repair processes, as its role in chromatin remodeling makes damaged DNA accessible to repair proteins (98–101). Our results suggest that the increased DNA damage caused by the dual agent treatment leads to an increased requirement of chromatin remodeling, and therefore increased interactions within the SWI / SNF complex. Others have shown that PARP1 physically interacts with members of the SWI / SNF complex in breast cancer cells (102), so PARP1 inhibition may directly impact SWI / SNF functions in DNA repair and transcription. TPCA also indicated a decreased interaction between p53 and tubulin with the combination treatment. Others have shown that p53 is directly associated with microtubules and is involved in centrosome duplication (91,92) and that inhibition of p53 localization to the mitotic centrosome results in breakdown of the centrosome and cell death (103). Taking this into consideration, the observed decreased interaction between p53 and tubulin could be due to altered microtubule dynamics resulting in paused cell cycle. We speculate that the accumulation of DNA damage from the dual agent treatment stalls the cell cycle in an attempt to repair DNA prior to division and ultimately leads to cell death.

Together, our results show that combination treatment of IB-DNQ and Rucaparib is a promising tumor-selective treatment strategy that targets cell cycle, DNA damage repair, and RNA Polymerase II transcription machinery only in cells with highly expressed NQO1. Future studies in mammalian systems and TNBC patient tumors are needed to better understand the efficacy and toxicity of this dual agent treatment in complex heterogeneous systems.

## METHODS

### Cell culture and drug treatment

Human MDA-MB-231 and MDA-MB-468 triple negative breast cancer cells were obtained from the American Tissue Culture Collection (ATCC, Manasas, VA). These cells natively express the NQO1*2 (609C>T) polymorphism rendering the protein unstable and shows no enzyme activity (104). The corresponding isogenic cell lines containing wildtype NQO1 were generated by the Boothman lab as previously described (18,105,106). Cells were grown in complete DMEM medium containing 5% FBS, and the isogenic cell lines containing pCDNA3 empty vector control and pCDNA3-NQO1 were intermittently selected in geneticin (400 μg/mL), and NQO1 expression were verified by Western blot. All cancer cell lines used in the study were tested for mycoplasma contamination routinely and grown in a 37 °C humidified incubator with 5% CO_2_, 95% air atmosphere.

NQO1 positive and NQO1*2 cells were treated with 0.1 μM IB-DNQ (2 hours), 0.4 μM IB-DNQ (2 hours). 15 μM Rucaparib (4 hours), or 0.1% DMSO (2 hours). For the combination treatment, cells were pretreated with 15 µM Rucaparib for 2 hours, then co-treated with 0.1 μM IB-DNQ and 15 μM Rucaparib for 2 hours. All drugs and DMSO were prepared in 5% DMEM.

### Relative Cell Survival Assays

Cells were pre-treated with Vehicle control (0.1% DMSO) or Rucaparib (15 µM) for two hours followed by treatment with various concentrations of IB-DNQ as indicated in combination with vehicle control (0.1% DMSO) or Rucaparib (15 µM) for additional two hours. Fresh media were added and allowed to grow until control cells became confluent (∼7 days). Relative long-term survival assays were based on assessment of cellular DNA content measured via Hoechst 33258 fluorescence signal (18) detected using VictorX3 Multimode Plate Reader. Data were expressed as percentage of relative surviving cells (% T/C, treatment/control) from at least three biological replicates and plotted using the GraphPad Prism software.

### Immunoblotting

Cells were lysed in cold RIPA buffer containing Halt^TM^ protease and phosphatase inhibitors (Thermo Scientific, Rockford, IL, USA) and processed using standard methods to collect the total cell lysate. Protein concentration of the whole-cell lysates was determined by BCA assay (Thermo Scientific, Waltham, MA, USA) and loading volume was normalized for all samples and ran on acrylamide SDS-PAGE gels, transferred to PVDF membrane, and processed using standard blotting techniques.

Primary antibodies for protein detection: NQO1 (A180, Cell Signaling Technology, Danvers, MA, USA), γH2AX (JBW301, MilliporeSigma, Burlington, MA, USA), PAR (10HA, Trevigen, Gaithersburg, MD, USA), and β-tubulin (D3U1W, Cell Signaling Technology, Danvers, MA, USA).

### Sample preparation for global and phosphoproteomics

Approximately 10 million cells were collected per treatment condition (n=3). Proteins were extracted by denaturing with 8M urea, followed by LysC / Trypsin Gold (Promega) digestion. Forty-five μg of protein was used for global analysis and 3 mg was used for phosphopeptide enrichment. Phosphopeptides were enriched by applying the 3 mg samples prepared above to High-Select TiO_2_ Phosphopeptide Enrichment tips (A32993, Thermo Fisher Scientific). Tips were washed and eluted using vendor-described protocols.

### TPP sample preparation

Approximately 10 million cells were collected per treatment condition (n=2). Cells were resuspended in NP-40 lysis buffer (40 mM HEPES (pH 7.5), 200 mM NaCl, 5 mM b-glycerophosphate, 0.1 mM sodium orthovanadate, Roche EDTA free complete protease inhibitor, 2 mM TCEP, 0.4% NP-40) and sonicated in a Bioruptor sonication system from Diagenode Inc. (30-s / 30-s on / off cycles for 30 min, 4°C). Following centrifugation at 12,000*g* for 15 min, protein concentrations were determined using a Bradford protein assay (5000002, Bio-Rad). Samples were adjusted to the same concentration and divided equally into 8 tubes for heat treatment in the thermocycler (25°C for 3 minutes, 3 minutes at temperature gradient, 25°C for 3 minutes).

Temperature gradient is as follows: 25°C, 35°C, 39.3°C, 50.1°C, 54.8°C, 60.7°C, 74.9°C, 90°C. Each sample was centrifuged to pellet aggregated protein, and supernatant was reserved. Proteins were precipitated in 20% trichloroacetic acid (TCA) overnight and washed with acetone. Precipitated proteins were resuspended in 8M Urea in 100 mM HCl (pH 8.0). Proteins were reduced with 5 mM tris(2-carboxyethyl)phosphine (TCEP) and alkylated with 10 mM chloroacetaminde. Samples incubated with Trypsin / Lys-C mix (1:70 protease:substrate ratio; V5072, Promega) for 4 hours, then were diluted with 100 mM Tris-HCl to reduce urea concentration to less than 1M and digested overnight. Digestion was quenched with 0.4% trifluoroacetic acid (TFA).

### Peptide purification, TMT-labeling, and high-pH basic fractionation

Peptides were desalted on a 50 mg Sep-Pak C18 Vac (Waters Corporation) using a vacuum manifold. Peptides were eluted with 70% acetonitrile (ACN) and 0.1% formic acid (FA), dried by speed vacuum, and resuspended in 50 mM triethylammonium bicarbonate (TEAB, pH 8.5). For global and phosphoproteomics samples, peptide concentration was measured using the Pierce Quantitative Colorimetric Peptide Assay Kit (23275, Thermo Fisher Scientific) to ensure an equal amount of each sample was labeled. Samples were then tandem mass tag (TMT)-labeled for 2 hours at room temperature. Labeling reaction was quenched with 5% hydroxylamine, and labeled peptides were mixed together. Peptide mixtures were desalted and fractionated using 50 mg Sep-Pak C18 Vac columns (Waters Corporation). Peptides were eluted in 12.5%, 15%, 17.5%, 20%, 22.5%, 25%, 30%, and 70% acetonitrile in 0.1% triethylamine.

Fractions were dried in speed vacuum and resuspended in 0.1% formic acid.

### Global proteomics data acquisition: Nano-LC-MS / MS

Nano-LC–MS / MS analyses were performed on an Orbitrap Fusion Lumos mass spectrometer (Thermo Fisher Scientific) coupled to an EASY-nLC HPLC system (Thermo Fisher Scientific). Each fraction was loaded onto a reversed-phase PepMap RSLC C18 column with an Easy-Spray tip at 400 nl / min (ES802A, Thermo Fisher Scientific; 2 μm, 100 Å, 75 μm by 25 cm). Peptides were eluted from 4 to 35% B over 160 min, 35 to 50% B for 14 min, and dropping from 50 to 10% B over the final 1 min (mobile phases A: 0.1% FA and water; B: 0.1% FA and 80% ACN). Mass spectrometer settings include a capillary temperature of 300°C, and ion spray voltage was kept at 1.8 kV. The mass spectrometer method was operated in positive-ion mode with a 4-s cycle time data-dependent acquisition with advanced peak determination and Easy-IC on (internal calibrant). Precursor scans [mass / charge ratio (*m* / *z*), 400 to 1500] were done with an Orbitrap resolution of 60,000, 30% RF (radio frequency) lens, 50-ms maximum inject time (IT) and automatic gain control (AGC) target of 400,000, including charges of 2 to 6 for fragmentation with 60-s dynamic exclusion. Higher-energy collisional dissociation (HCD) MS2 scans were performed at 50k Orbitrap resolution, fixed collision energy of 38%, AGC target of 10,000, and 86-ms maximum IT.

### Phosphoproteomics data acquisition: Nano-LC-MS / MS / MS

Nano-LC–MS / MS / MS analyses were performed on an Orbitrap Fusion Lumos mass spectrometer (Thermo Fisher Scientific) coupled to an EASY-nLC HPLC system (Thermo Fisher Scientific). Each fraction was analyzed with the same LC conditions as above on a Lumos Orbitrap mass spectrometer (Thermo Fisher Scientific). Mass spectrometer settings include a capillary temperature of 300°C, and ion spray voltage was kept at 1.9 kV. The mass spectrometer method was operated in positive-ion mode with a 3-s cycle time data-dependent acquisition with advanced peak determination and Easy-IC on. Precursor scans (*m / z*, 400 to 1500) were done with an Orbitrap resolution of 120,000, 30% RF lens, 50-ms maximum IT, AGC target of 400,000, including charges of 2 to 7 for fragmentation with 90-s dynamic exclusion. Collision induced dissociation (CID) MS2 scans were performed at 30k Orbitrap resolution, fixed collision energy of 35%, AGC target of 50,000, and 60-ms maximum IT. HCD MS3 scans were performed at 50k Orbitrap resolution, fixed collision energy of 65%, AGC target of 100,000, and 105-ms maximum IT.

### TPP data acquisition: Nano-LC-MS / MS

Nano-LC–MS / MS analyses were performed on an Orbitrap Exploris 480 mass spectrometer (Thermo Fisher Scientific) coupled to an EASY-nLC HPLC system (Thermo Fisher Scientific). Two technical replicates of each fraction were loaded onto a reversed-phase PepMap RSLC C18 column with an Easy-Spray tip at 400 nl / min (ES802A, Thermo Fisher Scientific; 2 μm, 100 Å, 75 μm by 25 cm). Peptides were eluted from 6 to 32% B over 180 min, 10 to 32% B for 14 min, and dropping from 50 to 10% B over the final 1 min (mobile phases A: 0.1% FA and water; B: 0.1% FA and 80% ACN). Mass spectrometer settings include a capillary temperature of 300°C, and ion spray voltage was kept at 1.9 kV. The mass spectrometer method was operated in positive-ion mode with a 1.3-s cycle time data-dependent acquisition with advanced peak determination and Easy-IC on (internal calibrant). FAIMS CVs cycled through -45, -55, and -70 V. Precursor scans [mass / charge ratio (*m* / *z*), 375 to 1500] were done with an Orbitrap resolution of 60,000, 40% RF lens, normalized AGC target of 200%, including charges of 2 to 8 for fragmentation with 60-s dynamic exclusion. Higher-energy collisional dissociation (HCD) MS2 scans were performed at 45k Orbitrap resolution, normalized collision energy of 32%, normalized AGC target of 200%.

### Protein identification and quantification – global proteomics

Resulting MS2 RAW files were analyzed in Proteome Discoverer 2.5 (Thermo Fisher Scientific) with FASTA databases including UniProt human sequences plus common contaminants such as proteolytic digestion enzymes. Quantification methods used isotopic impurity levels available from Thermo Fisher Scientific. SEQUEST HT searches were conducted with a maximum number of **t**wo missed cleavages, a precursor mass tolerance of 10 parts per million, and a fragment mass tolerance of 0.02 Da. Static modifications used for the search were (i) carbamidomethylation on cysteine (C) residues and (ii) TMT6plex labels on lysine (K) residues and the N termini of peptides.

Dynamic modifications used for the search were oxidation of methionine, acetylation of N termini, and N-terminal loss of methionine with and without acetylation. Percolator false discovery rate (FDR) was set to a strict setting of 0.01 and a relaxed setting of 0.05. Values from both unique and razor peptides were used for quantification. In the consensus workflow, peptides were normalized by total peptide amount with scaling to mixed sample control. Global proteomics data have been deposited with MassIVE, project accession: MSV000094442.

### Protein identification and quantification – phosphoproteomics

Resulting MS3 RAW files were analyzed in Proteome Discoverer 2.5 (Thermo Fisher Scientific) with FASTA databases including UniProt human sequences plus common contaminants. Quantification methods used isotopic impurity levels available from Thermo Fisher Scientific. SEQUEST HT searches were conducted with a maximum number of two missed cleavages, a precursor mass tolerance of 10 parts per million, and a fragment mass tolerance of 0.02 Da. Static modifications used for the search were (i) carbamidomethylation on cysteine (C) residues and (ii) TMT6plex labels on lysine (K) residues and the N termini of peptides. Dynamic modifications used for the search were phosphorylation of serine, threonine, and tyrosine, oxidation of methionine and tryptophan, acetylation of N termini, and N-terminal loss of methionine with and without acetylation. Percolator false discovery rate (FDR) was set to a strict setting of 0.01 and a relaxed setting of 0.05. Values from both unique and razor peptides were used for quantification. In the consensus workflow, SPS Mass Matches threshold was 0, and peptides were normalized by total peptide amount with scaling to a mixed sample control. Phosphoproteomics data have been deposited with MassIVE, project accession: MSV000094442.

### Protein identification and quantification – TPP

Resulting MS2 RAW files were analyzed in Proteome Discovererä 2.5 (Thermo Fisher Scientific) with FASTA databases including UniProt human sequences plus common contaminants. Quantification methods used isotopic impurity levels available from Thermo Fisher Scientific. SEQUEST HT searches were conducted with a maximum number of three missed cleavages, a precursor mass tolerance of 10 parts per million, and a fragment mass tolerance of 0.02 Da. Static modifications used for the search were (i) carbamidomethylation on cysteine (C) residues and (ii) TMTpro labels on lysine (K) residues and the N termini of peptides. Dynamic modifications used for the search were oxidation of methionine, acetylation of N termini, and N-terminal loss of methionine with and without acetylation. Percolator false discovery rate (FDR) was set to a strict setting of 0.01 and a relaxed setting of 0.05. Values from both unique and razor peptides were used for quantification. In the consensus workflow, peptides were normalized by total peptide amount with no scaling. No normalization was performed for TPP experiments. Thermal proteome profiling data have been deposited with MassIVE, project accession: MSV000094442.

### Analysis of TPP data

The InflectSSP R-package (v 1.6) in R Studio (R Studio for Mac, version 2022.12.0+353) was used for curve fitting, melt temperature, and melt shift analysis. Accession numbers, number of PSMs, number of unique peptides, and abundance values for each condition were divided into separate Excel files. InflectSSP parameters were set as follows: NControl = 2, NCondition = 2, PSM = 0, UP =0, CurveRsq = 0, PValMelt = 0.05, MeltLimit = 0, STRINGScore = 0.99. Differential thermal proximity coaggregation analysis was conducted using the R package *Rtpca* (86) with the normalized and corrected abundances from the InflectSSP output. Only melt curves with an R-squared value >= 0.7 were used. For a given protein, the abundance values of biological replicates were averaged for each temperature for each condition. First, complexes with co-aggregation were tested using the runTPCA function and the CORUM human protein complex annotations with number of sampling set for 100,000 for each condition. Protein complexes with a p-value less than 0.15 in at least one condition were further analyzed for differential thermal proximity coaggregation between the two conditions. Differential TPCA was conducted using the runDiffTPCA function where the protein-protein interaction annotation was derived from BioGRID. Protein pairs with a p-value ≤ 0.05 were considered to have differential thermal proximity coaggregation. The list of CDK1 targets was obtained from Uniprot as of May 5, 2023. Volcano plots were created using ggplot2. The melt curves of binary protein pairs with differential TPCA were graphed using the plotPPiProfiles functions from Rtpca. CORUM v.4 human protein complex annotations were downloaded from http://mips.helmholtz-muenchen.de/corum/ (107,108). The physical protein-protein interaction data were used from BioGRID v.4.4.210 downloaded from https://thebiogrid.org (109).

### Kinase Substrate Enrichment Analysis (KSEA) (**50,51**)

The KSEA web app was used for analysis of phosphoproteomics data. The algorithm takes an input file in *.csv* format consisting of a list of peptides, corresponding protein and gene names, phosphosites and associated fold change (FC) and p-values. The *p*-value cutoff and substrate count cutoff were set at 0.05 and 2, respectively. Kinase substrate relationships from both PhosphoSitePlus and NetworkIn were used.

### Gene Ontology and Protein Network Analysis

ShinyGo (version 0.80) was used to determine enrichment in phosphoproteomics data. Analyzed proteins contained phosphopeptides that were significantly changing (p ≤ 0.05) in combination treatment (NQO1+) compared to DMSO. A background list containing all detected proteins was provided. FDR cutoff was set to 0.05. The PANTHER overrepresentation test (version 18.0) was used for gene ontology analysis of proteins with phosphopeptides increasing with combination treatment and decreasing with 0.4 µM IB-DNQ. A Fisher’s exact test was used to calculate enrichment, and a false discovery rate was calculated. Protein-protein interaction networks were analyzed using STRING (version 12.0).

### Data visualization

Volcano plots and PCA plots were created in R Studio using ggplot2(110) (version 3.4.2). Upset plots were created in R studio using UpSetR (111). Heatmap was created in ProteomeDiscoverer™ 2.5 (Thermo Fisher Scientific) using scaled abundance of peptide groups, Euclidean distance, Ward linkage method, and scaling before clustering.

## Declarations

### Ethics approval and consent to participate

Not applicable.

### Consent for publication

Not applicable.

## Author’s contributions

A.M.R. – data acquisition, data analysis, writing and revision of manuscript

H.R.S.W. – data acquisition, data analysis, revision of manuscript

S.A.P.J. – writing and revision of manuscript

A.B.W. – data acquisition, data analysis

G.R. – data analysis

J.J.S. – data acquisition, revision of manuscript

N.S. – experimental design, sample preparation

P.H. – development of IB-DNQ

D.A.B. – experimental design

E.A.M. – experimental design, revision of manuscript

A.L.M. – experimental design, data analysis, revision of manuscript

## Acknowledgements

We thank Neil McCracken for support with data analysis using InflectSSP and Colton Starcher for assistance with cell culture. The mass spectrometry work performed in this work was done by the Indiana University School of Medicine Center for Proteome Analysis. Acquisition of the IUSM Center for Proteome Analysis’ instrumentation used for this project was provided by the Indiana University Precision Health Initiative. The Center for Proteome Analysis receives support from 3UL1TR002529 and P30CA082709.

## Footnotes

David Boothman is included as an author posthumously.

---

**Supplementary Figure 1:**
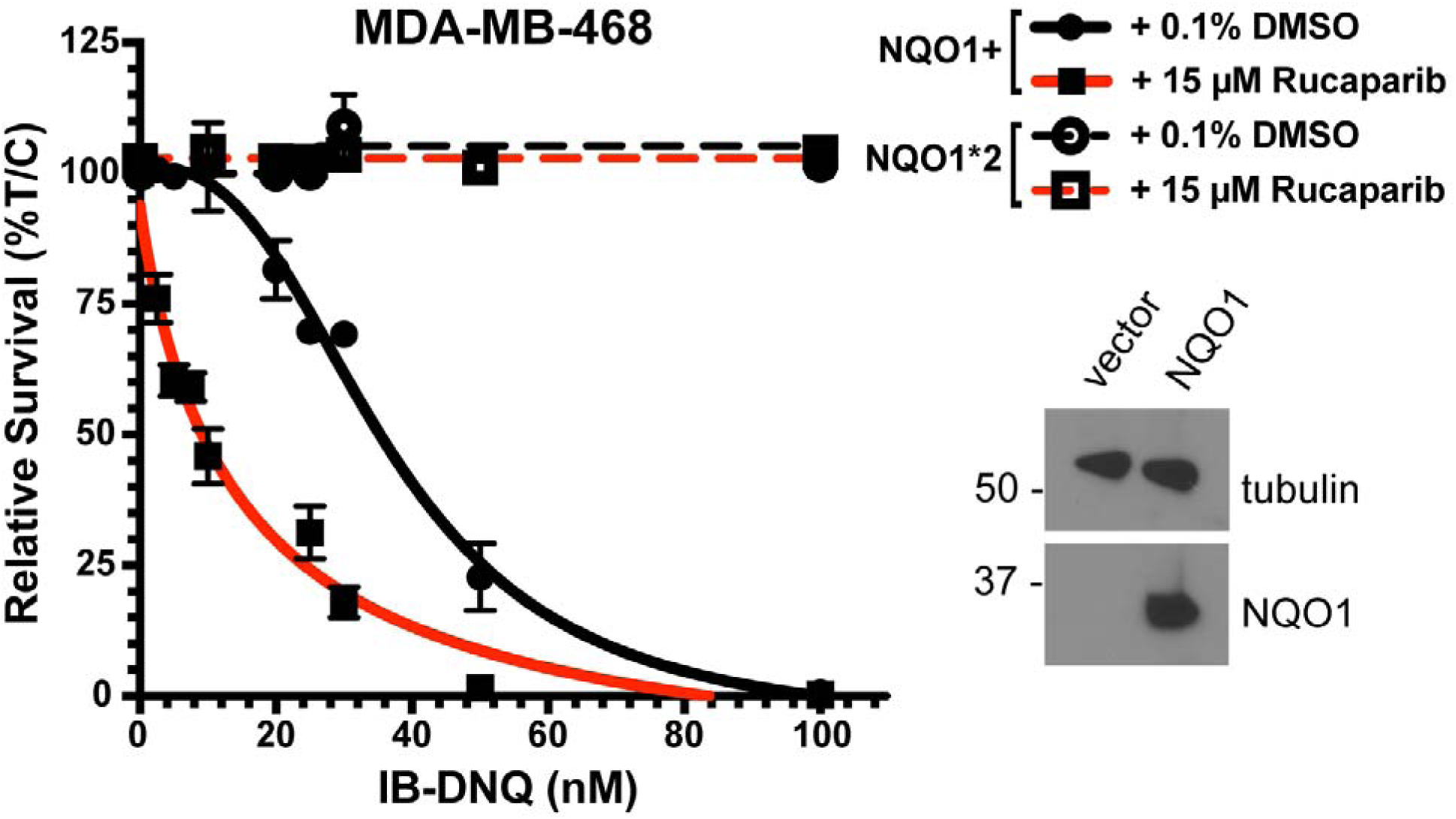
MDA-MB-468 triple negative breast cancer cells, which express the NQO1*2 variant (rapidly degraded NQO1), were altered with a vector to stably overexpress wild-type NQO1 (NQO1+) or an empty vector control. Cells were treated with increasing doses of IB-DNQ in combination with DMSO or 15 µM Rucaparib. Cell survival is plotted for increasing IB-DNQ concentrations. NQO1*2 cells are represented by dashed lines, and NQO1+ cells are represented by solid lines. Black lines show co-treatment with DMSO, and red lines show co-treatment with Rucaparib. %T/C = % of treated / control cells. Western blot insets show abundance of NQO1 in NQO1+ and vector containing MDA-MB-231 cells. Tubulin was used as the loading control.

**Supplementary Figure 2:**
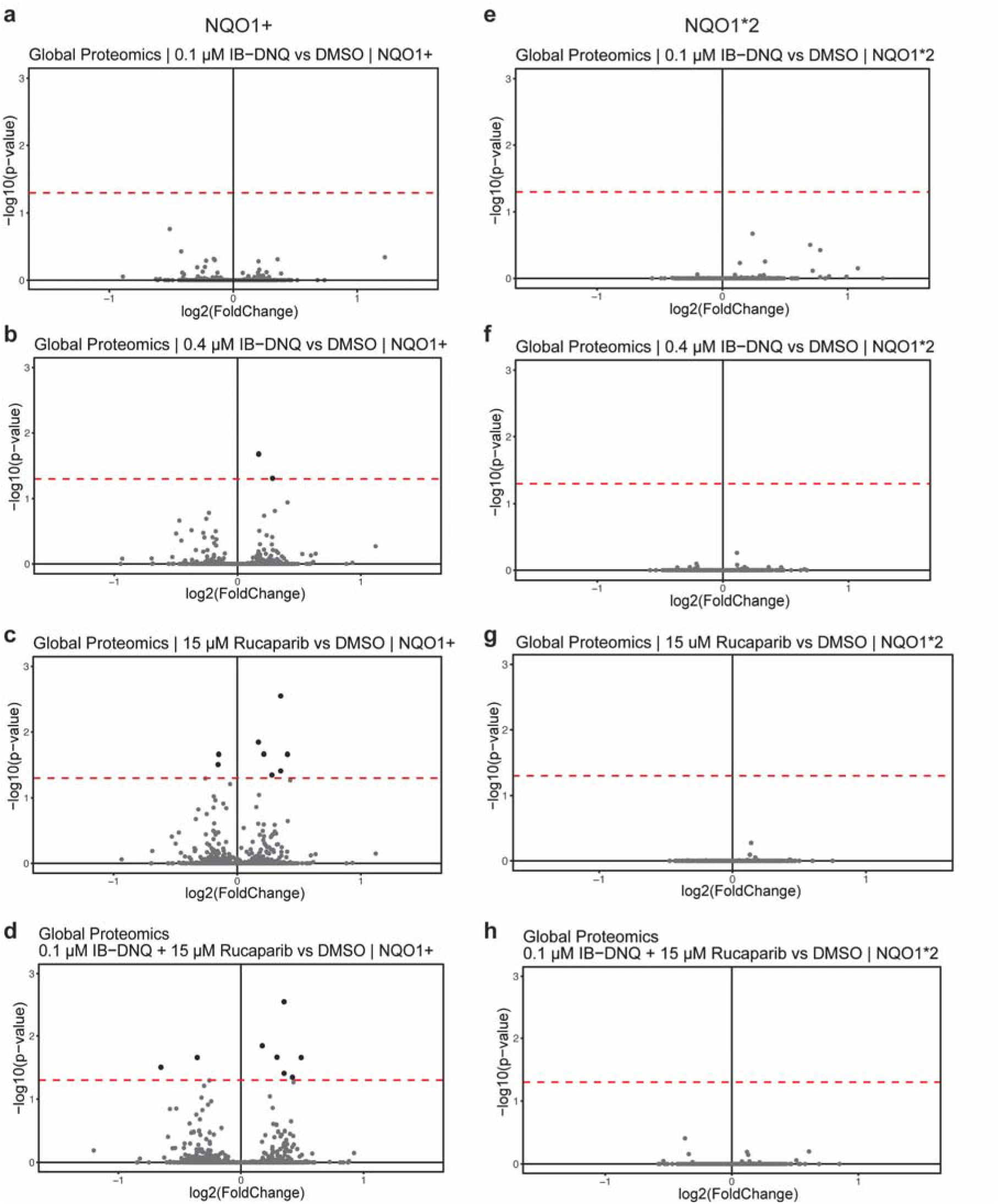
Global proteomics of NQO1+ (a-d) and NQO1*2 (e-h) MDA-MB-231 cells treated with 0.1 µM IB-DNQ (a,e), 0.4 µM IB-DNQ (b,f), 15 µM Rucaparib (c,g), or combination treatment (d,h). Red dashed line is p-value cut off p=0.05.

**Supplementary Figure 3:**
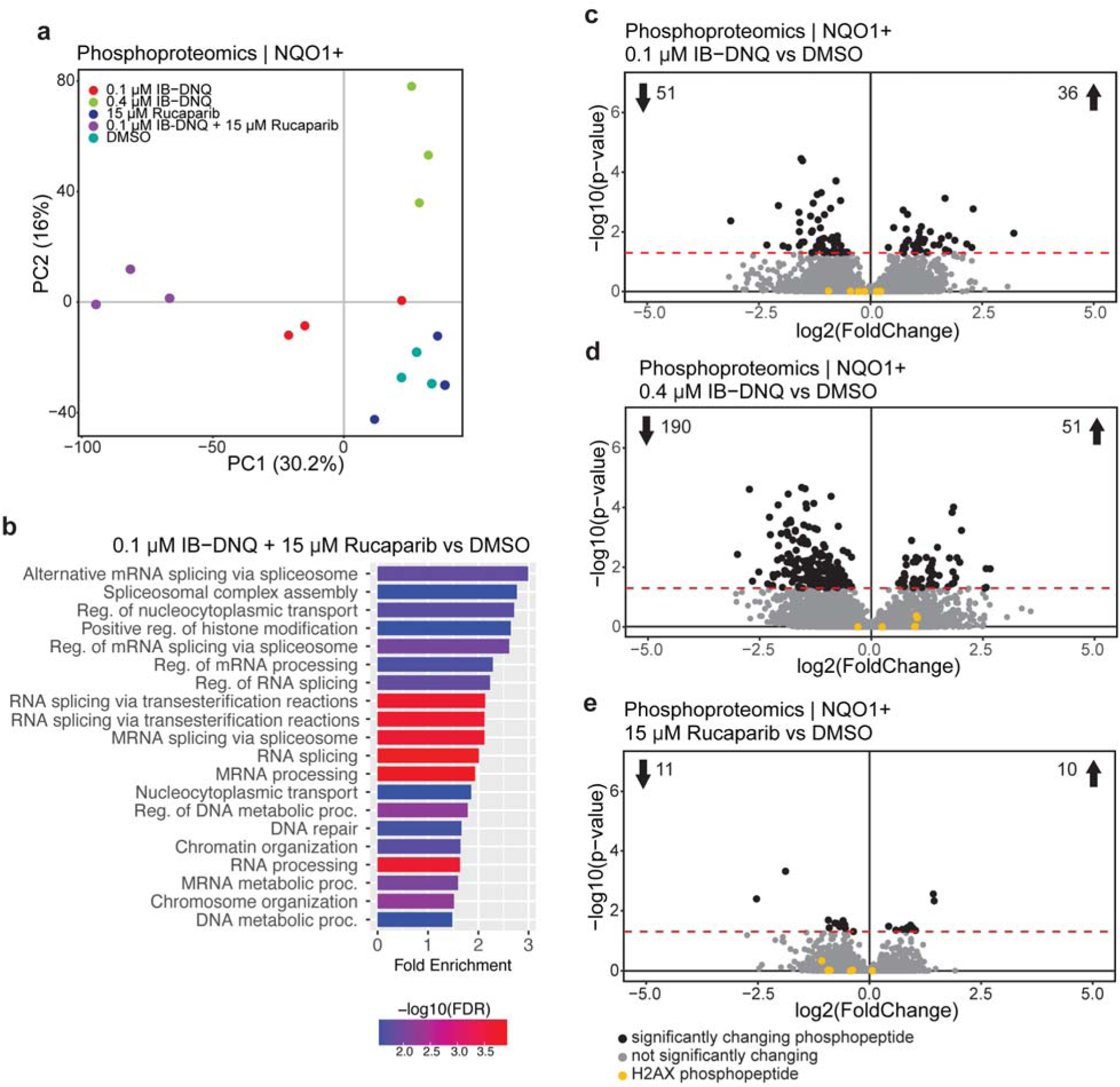
Changes in phosphoproteome are minimal in single agent treatment compared to combination treatment in NQO1+ cells. a) Principal component analysis of phosphoproteomics data from NQO1+ TNBC cells. Data was generated using scaled abundances of phosphopeptide groups that were quantified in all samples (n=3). Data from 3 independent replicates group together. Combination treated cells group together, separately from other treatment groups. b) Gene ontology analysis of proteins with significantly changing phosphopeptides in combination treatment compared to DMSO. c-e) Volcano plots show differential phosphorylation of phosphopeptide groups in 0.1 µM IB-DNQ (c), 0.4 µM IB-DNQ (d), and 15 µM Rucaparib (e), treated NQO1+ cells compared to DMSO treated cells. Yellow points represent phosphopeptides matching to the histone protein H2AX. Red dashed line is p-value cut off p=0.05.

**Supplementary Figure 4:**
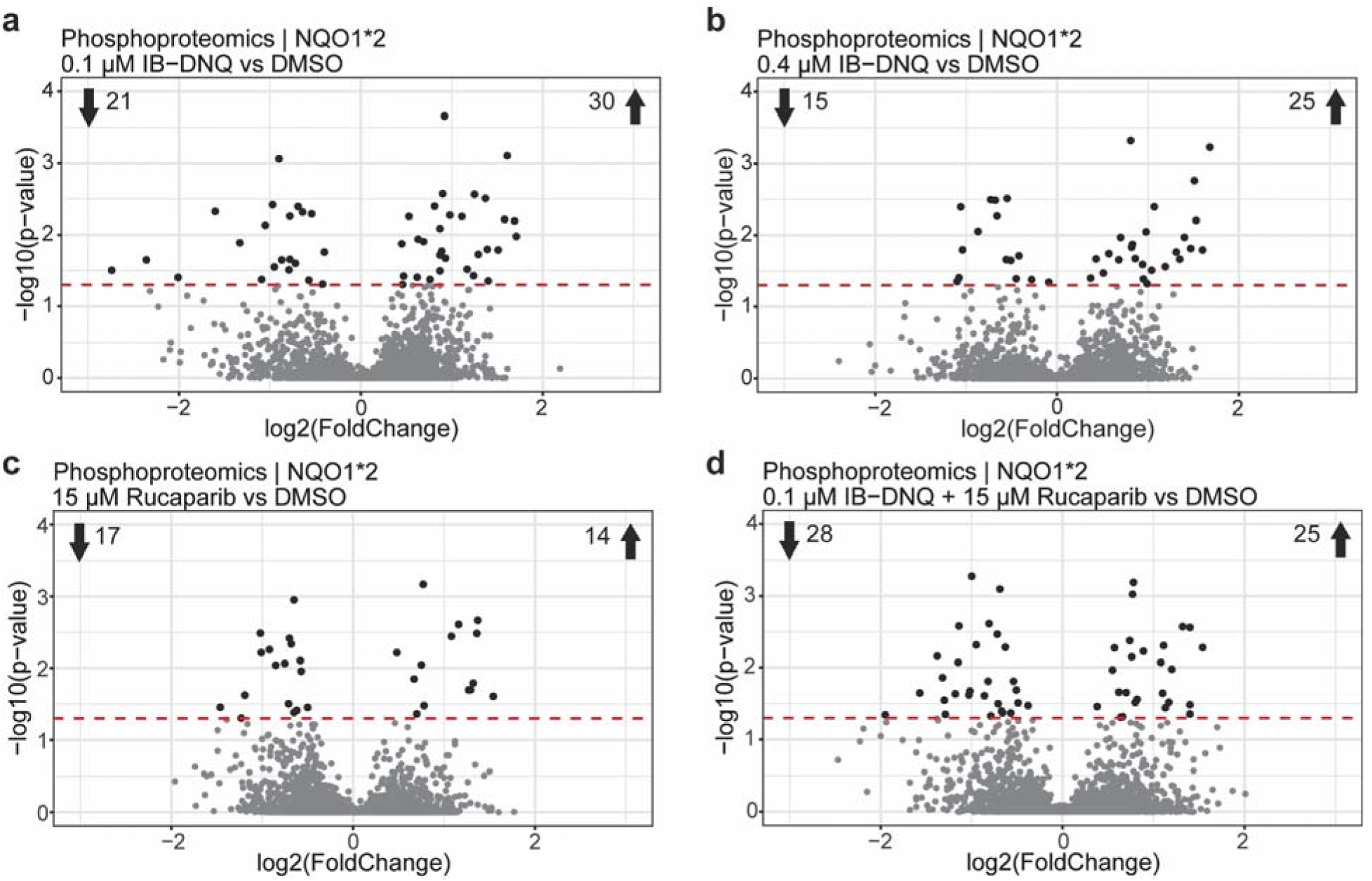
Phosphoproteomics of NQO1*2 MDA-MB-231 cells. Volcano plots show differential phosphorylation of peptide groups in drug treated NQO1*2 cells compared to DMSO treated cells. Red dashed line is p-value cut off p=0.05.

**Supplementary Figure 5:**
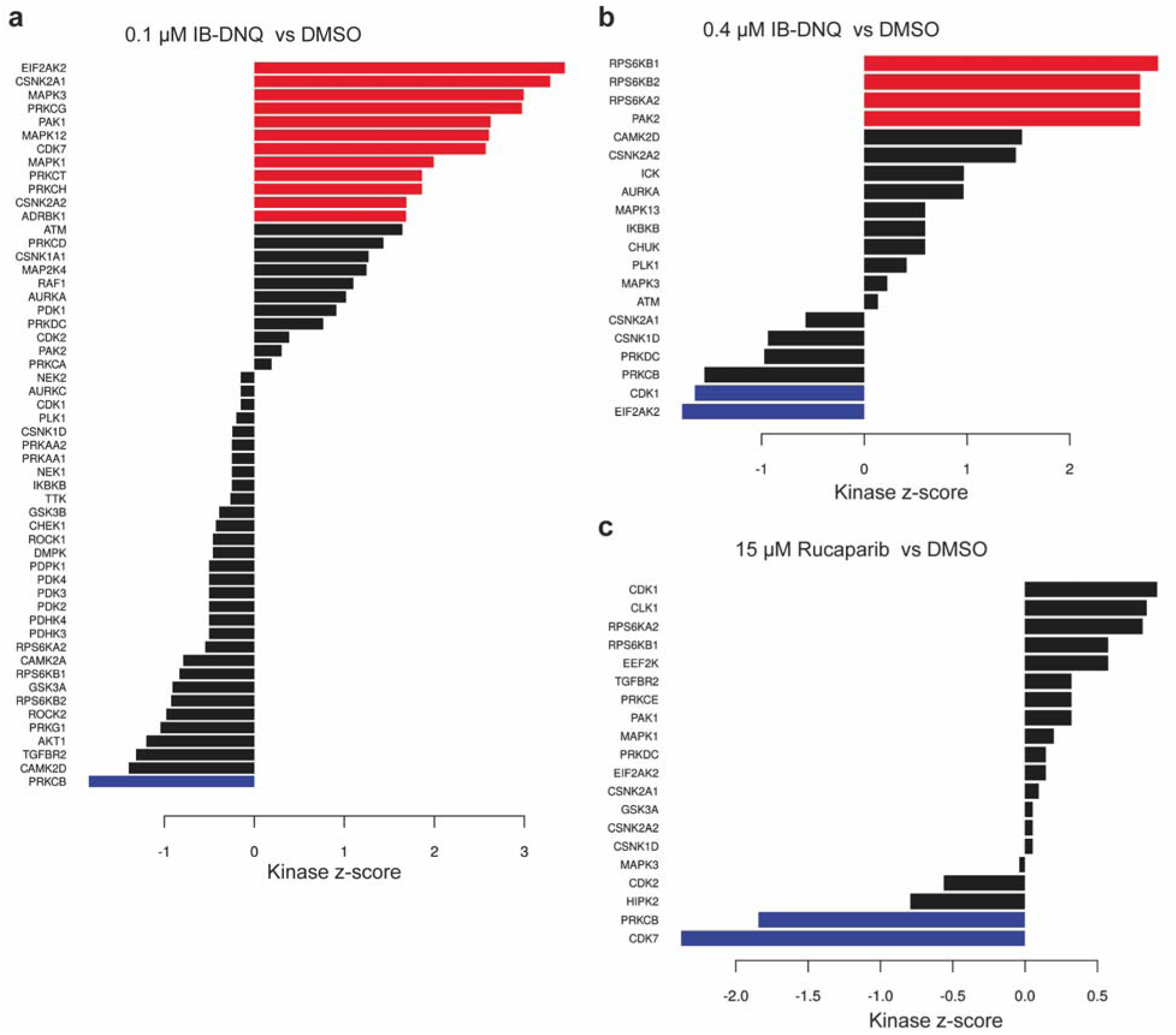
Kinase substrate enrichment analysis (KSEA) of NQO1+ single agent treated cells compared to DMSO. KSEA bar plots for 0.1 µM IB-DNQ (a), 0.4 µM IB-DNQ (b), and 15 µM Rucaparib (c). Red bars indicate kinases with significantly increased inferred activity (p ≤ 0.05), and blue bars indicate kinases with significantly decreased inferred activity. Black bars show kinases with differential inferred kinase activity but are not statistically significant.

**Supplementary Figure 6:**
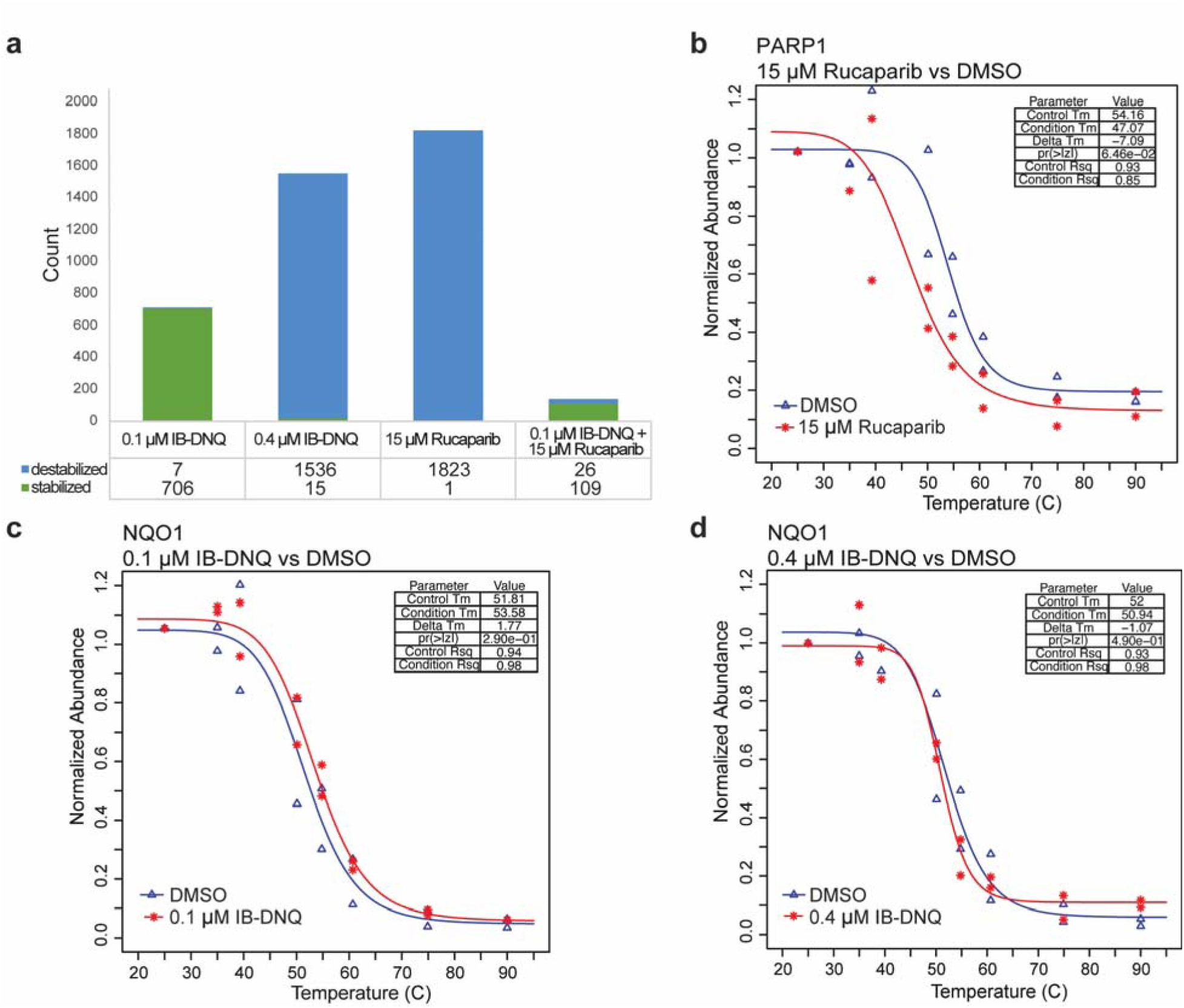
Thermal proteome profiling. a) Summary of stabilized and destabilized proteins in each condition compared to DMSO in NQO1+ cells. b) Melt curves for PARP1 (Rucaparib target) from 15 µM Rucaparib compared to DMSO treated NQO1+ cells. c,d) Melt curves for NQO1 (IB-DNQ target) from 0.1 µM IB-DNQ treated NQO1+ cells (c) and 0.4 µM IB-DNQ treated NQO1+ cells (d) compared to DMSO.

**Supplementary Figure 7:**
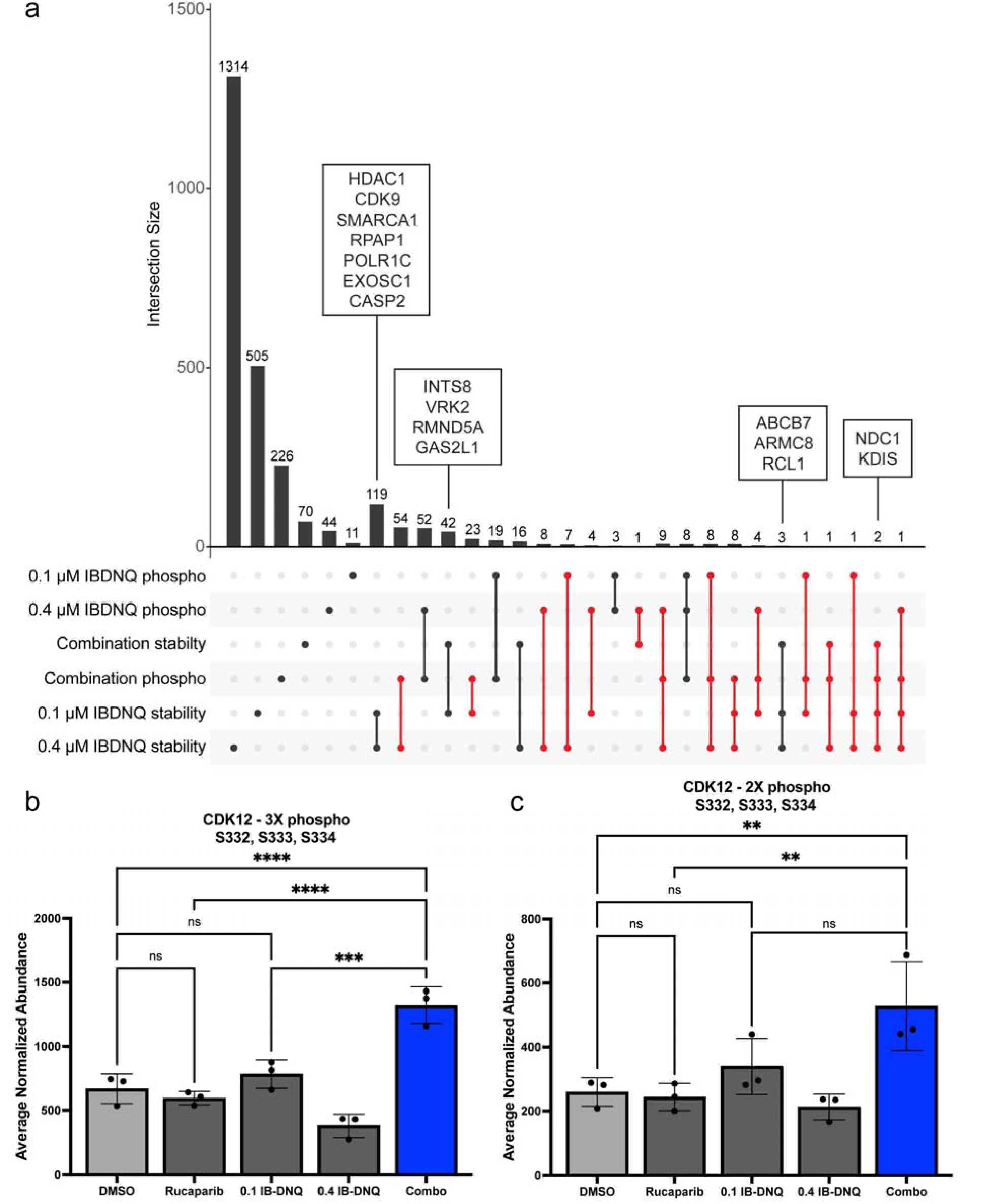
Comparison of proteins with phosphorylation and thermal stability changes in each of the IB-DNQ treatment groups (0.1 µM, 0.4 µM, and combination treatment). a) Upset plot showing overlap of phosphorylation and thermal stability changes. Intersection size is the number of unique or overlapping proteins or peptides. Single dot represents proteins or peptides that are unique to that category, and 2 or more dots connected with a line show where there are overlapping proteins. Red connections show overlaps of stability and phosphorylation changes. Boxes show some of the genes identified in certain categories. b,c) Average normalized abundance of phosphopeptides for CDK12 in each of the treatment groups. Phosphorylated residue is shown above each graph. b) Abundances for 3X phosphorylated peptide RRSSSPFLSK. c) Abundances for 2X phosphorylated peptide RRSSSPFLSK. Phosphorylation is ambiguous for serines 332, 333, and 334. Light gray bars show abundance in DMSO, dark gray bars show abundance in single agent treatments, and blue bars show abundance for the IB-DNQ and Rucaparib combination treatment. Comparisons between groups show statistical significance as calculated by one-way ANOVA and Fisher LSD multiple comparison test. ns = not significant, * = p<0.05, ** = p<0.01, *** = p<0.001, **** = p<0.0001.

## REFERENCES

1. Yin L, Duan JJ, Bian XW, Yu SC. Triple-negative breast cancer molecular subtyping and treatment progress. Breast Cancer Res BCR. 2020 Jun 9;22(1):61.

2. Almansour NM. Triple-Negative Breast Cancer: A Brief Review About Epidemiology, Risk Factors, Signaling Pathways, Treatment and Role of Artificial Intelligence. Front Mol Biosci. 2022 Jan 25;9:836417.

3. Howard FM, Olopade OI. Epidemiology of Triple-Negative Breast Cancer: A Review. Cancer J Sudbury Mass. 2021 Feb 1;27(1):8–16.

4. Brenton JD, Carey LA, Ahmed AA, Caldas C. Molecular Classification and Molecular Forecasting of Breast Cancer: Ready for Clinical Application? J Clin Oncol. 2005 Oct 10;23(29):7350–60.

5. Foulkes WD, Smith IE, Reis-Filho JS. Triple-Negative Breast Cancer. N Engl J Med. 2010 Nov 11;363(20):1938–48.

6. Motea EA, Huang X, Singh N, Kilgore J, Williams N, Xie XJ, et al. NQO1-dependent, tumor-selective radiosensitization of non-small cell lung cancers. Clin Cancer Res Off J Am Assoc Cancer Res. 2019 Apr 15;25(8):2601–9.

7. Huang X, Dong Y, Bey EA, Kilgore JA, Bair JS, Li LS, et al. An NQO1 Substrate with Potent Antitumor Activity That Selectively Kills by PARP1-Induced Programmed Necrosis. Cancer Res. 2012 Jun 14;72(12):3038–47.

8. Gerber DE, Beg MS, Fattah F, Frankel AE, Fatunde O, Arriaga Y, et al. Phase 1 study of ARQ 761, a β-lapachone analogue that promotes NQO1-mediated programmed cancer cell necrosis. Br J Cancer. 2018 Oct;119(8):928–36.

9. Parkinson EI, Bair JS, Cismesia M, Hergenrother PJ. Efficient NQO1 substrates are potent and selective anticancer agents. ACS Chem Biol. 2013 Oct 18;8(10):2173– 83.

10. Wettasinghe AP, Singh N, Starcher CL, DiTusa CC, Ishak-Boushaki Z, Kahanda D, et al. Detecting Attomolar DNA-Damaging Anticancer Drug Activity in Cell Lysates with Electrochemical DNA Devices. ACS Sens. 2021 Jul 23;6(7):2622–9.

11. Luo M, Shen N, Shang L, Fang Z, Xin Y, Ma Y, et al. Simultaneous Targeting of NQO1 and SOD1 Eradicates Breast Cancer Stem Cells via Mitochondrial Futile Redox Cycling. Cancer Res. 2024 Dec 16;84(24):4264–82.

12. Huang X, Motea EA, Moore ZR, Yao J, Dong Y, Chakrabarti G, et al. Leveraging an NQO1 Bioactivatable Drug for Tumor-Selective Use of Poly(ADP-ribose) Polymerase Inhibitors. Cancer Cell. 2016 Dec;30(6):940–52.

13. Bey EA, Bentle MS, Reinicke KE, Dong Y, Yang CR, Girard L, et al. An NQO1- and PARP-1-mediated cell death pathway induced in non-small-cell lung cancer cells by β-lapachone. Proc Natl Acad Sci. 2007 Jul 10;104(28):11832–7.

14. Chakrabarti G, Silvers MA, Ilcheva M, Liu Y, Moore ZR, Luo X, et al. Tumor-selective use of DNA base excision repair inhibition in pancreatic cancer using the NQO1 bioactivatable drug, β-lapachone. Sci Rep. 2015 Nov 25;5:17066.

15. Siegel D, Ross D. Immunodetection of NAD(P)H:quinone oxidoreductase 1 (NQO1) in human tissues. Free Radic Biol Med. 2000 Aug;29(3–4):246–53.

16. Begleiter A, Fourie J. Induction of NQO1 in cancer cells. Methods Enzymol. 2004;382:320–51.

17. Deller S, Macheroux P, Sollner S. Flavin-dependent quinone reductases. Cell Mol Life Sci. 2008 Jan 1;65(1):141–60.

18. Pink JJ, Planchon SM, Tagliarino C, Varnes ME, Siegel D, Boothman DA. NAD(P)H:Quinone oxidoreductase activity is the principal determinant of beta-lapachone cytotoxicity. J Biol Chem. 2000 Feb 25;275(8):5416–24.

19. Finkel T, Holbrook NJ. Oxidants, oxidative stress and the biology of ageing. Nature. 2000;408(6809):239–47.

20. Wang M, Chen S, Ao D. Targeting DNA repair pathway in cancer: Mechanisms and clinical application. MedComm. 2021;2(4):654–91.

21. Torgovnick A, Schumacher B. DNA repair mechanisms in cancer development and therapy. Front Genet. 2015 Apr 23;6:157.

22. Rose M, Burgess JT, O’Byrne K, Richard DJ, Bolderson E. PARP Inhibitors: Clinical Relevance, Mechanisms of Action and Tumor Resistance. Front Cell Dev Biol. 2020 Sep 9;8:564601.

23. Dantzer F, De La Rubia G, Ménissier-De Murcia J, Hostomsky Z, De Murcia G, Schreiber V. Base excision repair is impaired in mammalian cells lacking poly(ADP-ribose) polymerase-1. Biochemistry. 2000;39(25):7559–69.

24. Wang M, Wu W, Wu W, Rosidi B, Zhang L, Wang H, et al. PARP-1 and Ku compete for repair of DNA double strand breaks by distinct NHEJ pathways. Nucleic Acids Res. 2006 Nov 1;34(21):6170–82.

25. Han Y, Yu X, Li S, Tian Y, Liu C. New Perspectives for Resistance to PARP Inhibitors in Triple-Negative Breast Cancer. Front Oncol [Internet]. 2020 Nov 25 [cited 2025 Mar 20];10. Available from: https://www.frontiersin.org/journals/oncology/articles/10.3389/fonc.2020.578095/full

26. Helleday T, Petermann E, Lundin C, Hodgson B, Sharma RA. DNA repair pathways as targets for cancer therapy. Nat Rev Cancer. 2008 Mar;8(3):193–204.

27. Lee J m., Ledermann JA, Kohn EC. PARP Inhibitors for BRCA1/2 mutation-associated and BRCA-like malignancies. Ann Oncol. 2014 Jan 1;25(1):32–40.

28. Ernster L, Ljunggren M, Danielson L. Purification and some properties of a highly dicumarol-sensitive liver diaphorase. Biochem Biophys Res Commun. 1960 Feb 1;2(2):88–92.

29. Timson DJ. Dicoumarol: A Drug which Hits at Least Two Very Different Targets in Vitamin K Metabolism. Curr Drug Targets. 2017;18(5):500–10.

30. Siegel D, Anwar A, Winski SL, Kepa JK, Zolman KL, Ross D. Rapid Polyubiquitination and Proteasomal Degradation of a Mutant Form of NAD(P)H:Quinone Oxidoreductase 1. Mol Pharmacol. 2001 Feb 1;59(2):263–8.

31. Gong P, Guo Z, Wang S, Gao S, Cao Q. Histone Phosphorylation in DNA Damage Response. Int J Mol Sci. 2025 Mar 7;26(6):2405.

32. Day M, Oliver AW, Pearl LH. Phosphorylation-dependent assembly of DNA damage response systems and the central roles of TOPBP1. DNA Repair. 2021 Dec;108:103232.

33. Keung MY, Wu Y, Badar F, Vadgama JV. Response of Breast Cancer Cells to PARP Inhibitors Is Independent of BRCA Status. J Clin Med. 2020 Mar 30;9(4):940.

34. Mah LJ, El-Osta A, Karagiannis TC. γH2AX: a sensitive molecular marker of DNA damage and repair. Leukemia. 2010 Apr;24(4):679–86.

35. Podhorecka M, Skladanowski A, Bozko P. H2AX Phosphorylation: Its Role in DNA Damage Response and Cancer Therapy. J Nucleic Acids. 2010 Aug 3;2010:920161.

36. Burma S, Chen BP, Murphy M, Kurimasa A, Chen DJ. ATM phosphorylates histone H2AX in response to DNA double-strand breaks. J Biol Chem. 2001 Nov 9;276(45):42462–7.

37. Ward IM, Chen J. Histone H2AX is phosphorylated in an ATR-dependent manner in response to replicational stress. J Biol Chem. 2001 Dec 21;276(51):47759–62.

38. Tholen J, Razew M, Weis F, Galej WP. Structural basis of branch site recognition by the human spliceosome. Science. 2022 Jan 7;375(6576):50–7.

39. Zhang Z, Will CL, Bertram K, Dybkov O, Hartmuth K, Agafonov DE, et al. Molecular architecture of the human 17S U2 snRNP. Nature. 2020 Jul;583(7815):310–3.

40. Loerch S, Leach JR, Horner SW, Maji D, Jenkins JL, Pulvino MJ, et al. The pre-mRNA splicing and transcription factor Tat-SF1 is a functional partner of the spliceosome SF3b1 subunit via a U2AF homology motif interface. J Biol Chem. 2019 Feb 22;294(8):2892–902.

41. Zhao J, Tian S, Guo Q, Bao K, Yu G, Wang X, et al. A PARylation-phosphorylation cascade promotes TOPBP1 loading and RPA-RAD51 exchange in homologous recombination. Mol Cell. 2022 Jul 21;82(14):2571–2587.e9.

42. Whitehouse CJ, Taylor RM, Thistlethwaite A, Zhang H, Karimi-Busheri F, Lasko DD, et al. XRCC1 stimulates human polynucleotide kinase activity at damaged DNA termini and accelerates DNA single-strand break repair. Cell. 2001 Jan 12;104(1):107–17.

43. Hoch NC, Hanzlikova H, Rulten SL, Tétreault M, Komulainen E, Ju L, et al. XRCC1 mutation is associated with PARP1 hyperactivation and cerebellar ataxia. Nature. 2017 Jan 5;541(7635):87–91.

44. Adamowicz M, Hailstone R, Demin AA, Komulainen E, Hanzlikova H, Brazina J, et al. XRCC1 protects transcription from toxic PARP1 activity during DNA base excision repair. Nat Cell Biol. 2021 Dec;23(12):1287–98.

45. Gottlieb TM, Jackson SP. The DNA-dependent protein kinase: requirement for DNA ends and association with Ku antigen. Cell. 1993 Jan 15;72(1):131–42.

46. Smith GCM, Jackson SP. The DNA-dependent protein kinase. Genes Dev. 1999 Apr 15;13(8):916–34.

47. Wechsler T, Chen BPC, Harper R, Morotomi-Yano K, Huang BCB, Meek K, et al. DNA-PKcs function regulated specifically by protein phosphatase 5. Proc Natl Acad Sci U S A. 2004 Feb 3;101(5):1247–52.

48. Chen S, Lee L, Naila T, Fishbain S, Wang A, Tomkinson AE, et al. Structural basis of Long-range to Short-range synaptic transition in NHEJ. Nature. 2021 May;593(7858):294–8.

49. Ma Y, Pannicke U, Schwarz K, Lieber MR. Hairpin opening and overhang processing by an Artemis/DNA-dependent protein kinase complex in nonhomologous end joining and V(D)J recombination. Cell. 2002 Mar 22;108(6):781–94.

50. Wiredja DD, Koyutürk M, Chance MR. The KSEA App: a web-based tool for kinase activity inference from quantitative phosphoproteomics. Bioinformatics. 2017 Nov 1;33(21):3489–91.

51. Casado P, Rodriguez-Prados JC, Cosulich SC, Guichard S, Vanhaesebroeck B, Joel S, et al. Kinase-substrate enrichment analysis provides insights into the heterogeneity of signaling pathway activation in leukemia cells. Sci Signal. 2013 Mar 26;6(268):rs6.

52. Hornbeck PV, Zhang B, Murray B, Kornhauser JM, Latham V, Skrzypek E. PhosphoSitePlus, 2014: mutations, PTMs and recalibrations. Nucleic Acids Res. 2015 Jan 28;43(D1):D512–20.

53. Horn H, Schoof EM, Kim J, Robin X, Miller ML, Diella F, et al. KinomeXplorer: an integrated platform for kinome biology studies. Nat Methods. 2014 Jun;11(6):603–4.

54. Linding R, Jensen LJ, Ostheimer GJ, van Vugt MATM, Jørgensen C, Miron IM, et al. Systematic discovery of in vivo phosphorylation networks. Cell. 2007 Jun 29;129(7):1415–26.

55. Linding R, Jensen LJ, Pasculescu A, Olhovsky M, Colwill K, Bork P, et al. NetworKIN: a resource for exploring cellular phosphorylation networks. Nucleic Acids Res. 2008 Jan;36(Database issue):D695–699.

56. Matthews HK, Bertoli C, de Bruin RAM. Cell cycle control in cancer. Nat Rev Mol Cell Biol. 2022 Jan;23(1):74–88.

57. Johnson N, Shapiro GI. Cyclin-dependent kinases (cdks) and the DNA damage response: rationale for cdk inhibitor–chemotherapy combinations as an anticancer strategy for solid tumors. Expert Opin Ther Targets. 2010 Nov;14(11):1199–212.

58. Fang Y, Zhang X. Targeting NEK2 as a promising therapeutic approach for cancer treatment. Cell Cycle. 2016 Mar 28;15(7):895–907.

59. Sasai K, Katayama H, Stenoien DL, Fujii S, Honda R, Kimura M, et al. Aurora-C kinase is a novel chromosomal passenger protein that can complement Aurora-B kinase function in mitotic cells. Cell Motil Cytoskeleton. 2004 Dec;59(4):249–63.

60. Bignone PA, Lee KY, Liu Y, Emilion G, Finch J, Soosay AER, et al. RPS6KA2, a putative tumour suppressor gene at 6q27 in sporadic epithelial ovarian cancer. Oncogene. 2007 Feb 1;26(5):683–700.

61. Mitsushima M, Toyoshima F, Nishida E. Dual role of Cdc42 in spindle orientation control of adherent cells. Mol Cell Biol. 2009 May;29(10):2816–27.

62. Archambault V, Lépine G, Kachaner D. Understanding the Polo Kinase machine. Oncogene. 2015 Sep;34(37):4799–807.

63. Reinhardt HC, Yaffe MB. Kinases that Control the Cell Cycle in Response to DNA Damage: Chk1, Chk2, and MK2. Curr Opin Cell Biol. 2009 Apr;21(2):245–55.

64. Songyang Z, Blechner S, Hoagland N, Hoekstra MF, Piwnica-Worms H, Cantley LC. Use of an oriented peptide library to determine the optimal substrates of protein kinases. Curr Biol. 1994 Nov 1;4(11):973–82.

65. Higashi H, Suzukitakahashi I, Taya Y, Segawa K, Nishimura S, Kitagawa M. Differences in Substrate Specificity between Cdk2-Cyclin A and Cdk2-Cyclin E *in Vitro*. Biochem Biophys Res Commun. 1995 Nov 13;216(2):520–5.

66. Holmes JK, Solomon MJ. A Predictive Scale for Evaluating Cyclin-dependent Kinase Substrates: A COMPARISON OF p34cdc2 AND p33cdk2*. J Biol Chem. 1996 Oct 11;271(41):25240–6.

67. Kitagawa M, Higashi H, Jung HK, Suzuki-Takahashi I, Ikeda M, Tamai K, et al. The consensus motif for phosphorylation by cyclin D1-Cdk4 is different from that for phosphorylation by cyclin A/E-Cdk2. EMBO J. 1996 Dec;15(24):7060–9.

68. Zarkowska T, U S, Harlow E, Mittnacht S. Monoclonal antibodies specific for underphosphorylated retinoblastoma protein identify a cell cycle regulated phosphorylation site targeted by CDKs. Oncogene. 1997 Jan;14(2):249–54.

69. Hsin JP, Manley JL. The RNA polymerase II CTD coordinates transcription and RNA processing. Genes Dev. 2012 Oct 1;26(19):2119–37.

70. Jeronimo C, Forget D, Bouchard A, Li Q, Chua G, Poitras C, et al. Systematic analysis of the protein interaction network for the human transcription machinery reveals the identity of the 7SK capping enzyme. Mol Cell. 2007 Jul 20;27(2):262–74.

71. Brackertz M, Gong Z, Leers J, Renkawitz R. p66alpha and p66beta of the Mi-2/NuRD complex mediate MBD2 and histone interaction. Nucleic Acids Res. 2006;34(2):397–406.

72. Brackertz M, Boeke J, Zhang R, Renkawitz R. Two highly related p66 proteins comprise a new family of potent transcriptional repressors interacting with MBD2 and MBD3. J Biol Chem. 2002 Oct 25;277(43):40958–66.

73. Le Guezennec X, Vermeulen M, Brinkman AB, Hoeijmakers WAM, Cohen A, Lasonder E, et al. MBD2/NuRD and MBD3/NuRD, Two Distinct Complexes with Different Biochemical and Functional Properties. Mol Cell Biol. 2006 Feb;26(3):843– 51.

74. Spruijt CG, Luijsterburg MS, Menafra R, Lindeboom RGH, Jansen PWTC, Edupuganti RR, et al. ZMYND8 Co-localizes with NuRD on Target Genes and Regulates Poly(ADP-Ribose)-Dependent Recruitment of GATAD2A/NuRD to Sites of DNA Damage. Cell Rep. 2016 Oct 11;17(3):783–98.

75. Tan CSH, Go KD, Bisteau X, Dai L, Yong CH, Prabhu N, et al. Thermal proximity coaggregation for system-wide profiling of protein complex dynamics in cells. Science. 2018 Mar 9;359(6380):1170–7.

76. Peck Justice SA, Barron MP, Qi GD, Wijeratne HRS, Victorino JF, Simpson ER, et al. Mutant thermal proteome profiling for characterization of missense protein variants and their associated phenotypes within the proteome. J Biol Chem. 2020 Nov;295(48):16219–38.

77. Huang JX, Lee G, Cavanaugh KE, Chang JW, Gardel ML, Moellering RE. High throughput discovery of functional protein modifications by Hotspot Thermal Profiling. Nat Methods. 2019 Sep;16(9):894–901.

78. Becher I, Andrés-Pons A, Romanov N, Stein F, Schramm M, Baudin F, et al. Pervasive Protein Thermal Stability Variation during the Cell Cycle. Cell. 2018 May 31;173(6):1495–1507.e18.

79. Martinez Molina D, Jafari R, Ignatushchenko M, Seki T, Larsson EA, Dan C, et al. Monitoring drug target engagement in cells and tissues using the cellular thermal shift assay. Science. 2013 Jul 5;341(6141):84–7.

80. McCracken NA, Peck Justice SA, Wijeratne AB, Mosley AL. Inflect: Optimizing Computational Workflows for Thermal Proteome Profiling Data Analysis. J Proteome Res. 2021 Apr 2;20(4):1874–88.

81. McCracken NA, Liu H, Runnebohm AM, Wijeratne HRS, Wijeratne AB, Staschke KA, et al. Obtaining Functional Proteomics Insights From Thermal Proteome Profiling Through Optimized Melt Shift Calculation and Statistical Analysis With InflectSSP. Mol Cell Proteomics MCP. 2023 Aug 9;22(9):100630.

82. Reed TJ, Tyl MD, Tadych A, Troyanskaya OG, Cristea IM. Tapioca: a platform for predicting de novo protein-protein interactions in dynamic contexts. Nat Methods. 2024 Mar;21(3):488–500.

83. Sharma D, De Falco L, Padavattan S, Rao C, Geifman-Shochat S, Liu CF, et al. PARP1 exhibits enhanced association and catalytic efficiency with γH2A.X-nucleosome. Nat Commun. 2019 Dec;10(1):5751.

84. Clark NJ, Kramer M, Muthurajan UM, Luger K. Alternative modes of binding of poly(ADP-ribose) polymerase 1 to free DNA and nucleosomes. J Biol Chem. 2012 Sep 21;287(39):32430–9.

85. Sun S, Zheng Z, Wang J, Li F, He A, Lai K, et al. Improved in situ characterization of protein complex dynamics at scale with thermal proximity co-aggregation. Nat Commun. 2023 Nov 24;14(1):7697.

86. Kurzawa N, Mateus A, Savitski MM. Rtpca: an R package for differential thermal proximity coaggregation analysis. Luigi Martelli P, editor. Bioinformatics. 2021 Apr 20;37(3):431–3.

87. Mathur R, Roberts CWM. SWI/SNF (BAF) Complexes: Guardians of the Epigenome. Annu Rev Cancer Biol. 2018;2(1):413–27.

88. Breitkreutz A, Choi H, Sharom JR, Boucher L, Neduva V, Larsen B, et al. A Global Protein Kinase and Phosphatase Interaction Network in Yeast. Science. 2010 May 21;328(5981):1043–6.

89. Rega C, Tsitsa I, Roumeliotis TI, Krystkowiak I, Portillo M, Yu L, et al. High resolution profiling of cell cycle-dependent protein and phosphorylation abundance changes in non-transformed cells. Nat Commun. 2025 Mar 16;16(1):2579.

90. Wu C, Ba Q, Lu D, Li W, Salovska B, Hou P, et al. Global and Site-Specific Effect of Phosphorylation on Protein Turnover. Dev Cell. 2021 Jan 11;56(1):111–124.e6.

91. Tarapore P, Horn HF, Tokuyama Y, Fukasawa K. Direct regulation of the centrosome duplication cycle by the p53-p21Waf1/Cip1 pathway. Oncogene. 2001 May;20(25):3173–84.

92. Giannakakou P, Sackett DL, Ward Y, Webster KR, Blagosklonny MV, Fojo T. p53 is associated with cellular microtubules and is transported to the nucleus by dynein. Nat Cell Biol. 2000 Oct;2(10):709–17.

93. Lin Z, Tan C, Qiu Q, Kong S, Yang H, Zhao F, et al. Ubiquitin-specific protease 22 is a deubiquitinase of CCNB1. Cell Discov. 2015;1:15028-.

94. Dubbury SJ, Boutz PL, Sharp PA. CDK12 regulates DNA repair genes by suppressing intronic polyadenylation. Nature. 2018 Dec;564(7734):141–5.

95. Potel CM, Kurzawa N, Becher I, Typas A, Mateus A, Savitski MM. Impact of phosphorylation on thermal stability of proteins. Nat Methods. 2021 Jul;18(7):757–9.

96. Smith IR, Hess KN, Bakhtina AA, Valente AS, Rodríguez-Mias RA, Villén J. Identification of phosphosites that alter protein thermal stability. Nat Methods. 2021 Jul;18(7):760–2.

97. Viktorsson K, Rieckmann T, Fleischmann M, Diefenhardt M, Hehlgans S, Rödel F. Advances in molecular targeted therapies to increase efficacy of (chemo)radiation therapy. Strahlenther Onkol. 2023;199(12):1091–109.

98. Ribeiro-Silva C, Vermeulen W, Lans H. SWI/SNF: Complex complexes in genome stability and cancer. DNA Repair. 2019 May;77:87–95.

99. Harrod A, Lane KA, Downs JA. The role of the SWI/SNF chromatin remodelling complex in the response to DNA double strand breaks. DNA Repair. 2020 Sep;93:102919.

100. Lans H, Marteijn JA, Vermeulen W. ATP-dependent chromatin remodeling in the DNA-damage response. Epigenetics Chromatin. 2012 Jan 30;5(1):4.

101. Davó-Martínez C, Helfricht A, Ribeiro-Silva C, Raams A, Tresini M, Uruci S, et al. Different SWI/SNF complexes coordinately promote R-loop- and RAD52-dependent transcription-coupled homologous recombination. Nucleic Acids Res. 2023 Jul 20;51(17):9055–74.

102. Sobczak M, Pitt AR, Spickett CM, Robaszkiewicz A. PARP1 Co-Regulates EP300– BRG1-Dependent Transcription of Genes Involved in Breast Cancer Cell Proliferation and DNA Repair. Cancers. 2019 Oct 11;11(10):1539.

103. Contadini C, Monteonofrio L, Virdia I, Prodosmo A, Valente D, Chessa L, et al. p53 mitotic centrosome localization preserves centrosome integrity and works as sensor for the mitotic surveillance pathway. Cell Death Dis. 2019 Nov 7;10(11):1–16.

104. Ross D, Kepa JK, Winski SL, Beall HD, Anwar A, Siegel D. NAD(P)H:quinone oxidoreductase 1 (NQO1): chemoprotection, bioactivation, gene regulation and genetic polymorphisms. Chem Biol Interact. 2000 Dec 1;129(1–2):77–97.

105. Cao L, Li LS, Spruell C, Xiao L, Chakrabarti G, Bey EA, et al. Tumor-selective, futile redox cycle-induced bystander effects elicited by NQO1 bioactivatable radiosensitizing drugs in triple-negative breast cancers. Antioxid Redox Signal. 2014 Jul 10;21(2):237–50.

106. Reinicke KE, Bey EA, Bentle MS, Pink JJ, Ingalls ST, Hoppel CL, et al. Development of beta-lapachone prodrugs for therapy against human cancer cells with elevated NAD(P)H:quinone oxidoreductase 1 levels. Clin Cancer Res Off J Am Assoc Cancer Res. 2005 Apr 15;11(8):3055–64.

107. Tsitsiridis G, Steinkamp R, Giurgiu M, Brauner B, Fobo G, Frishman G, et al. CORUM: the comprehensive resource of mammalian protein complexes–2022. Nucleic Acids Res. 2023 Jan 6;51(D1):D539–45.

108. Ruepp A, Brauner B, Dunger-Kaltenbach I, Frishman G, Montrone C, Stransky M, et al. CORUM: the comprehensive resource of mammalian protein complexes. Nucleic Acids Res. 2008 Jan 1;36(suppl_1):D646–50.

109. Stark C, Breitkreutz BJ, Reguly T, Boucher L, Breitkreutz A, Tyers M. BioGRID: a general repository for interaction datasets. Nucleic Acids Res. 2006 Jan 1;34(suppl_1):D535–9.

110. Wickham H. ggplot2: Elegant Graphics for Data Analysis [Internet]. New York, NY: Springer; 2009 [cited 2023 Jun 6]. Available from: https://link.springer.com/10.1007/978-0-387-98141-3

111. Conway JR, Lex A, Gehlenborg N. UpSetR: an R package for the visualization of intersecting sets and their properties. Bioinformatics. 2017 Sep 15;33(18):2938–40.

